# Minimizing co-growth as a broad predictor of community robustness

**DOI:** 10.64898/2026.04.14.717098

**Authors:** Milena S. Chakraverti-Wuerthwein, Yoshiya J. Matsubara, Finnegan D. Roach, Avaneesh V. Narla, Terence Hwa, Arvind Murugan

## Abstract

Microbial communities rarely remain in a fixed physiological state. Instead, they progress through internal life cycles in which changing metabolites, spatial organization, and physiological states reshape ecological interactions over time. Despite extensive theory on coexistence with fixed interactions, we lack simple quantitative predictors of robustness for communities undergoing repeated growth and dispersal cycles. Here we show that a single quantity, the temporal co-growth of community members, predicts robustness across several models of community maturation, including chemotactic spatial patterning, cross-feeding with toxicity, and a phenomenological many-species model with prescribed growth trajectories. Communities in which different species grow at distinct times persist far longer under stochastic reseeding than communities with overlapping growth, with average community lifetime increasing approximately exponentially as co-growth decreases. Across the systems studied here, diverse mechanisms such as spatial organization, metabolic cascades, and physiological programs promote robustness insofar as they reduce the temporal overlap of rapid growth across species. These results identify co-growth as a common quantitative feature of robust dynamically maturing communities and suggest that minimizing co-growth may provide a broader organizing principle for ecological robustness.

---

Microbial ecosystems are remarkable in their ability to grow and spread, while maintaining a stable species composition over long times despite environmental fluctuations. Traditionally, studies of community stability and robustness have focused on static environments, where microbial dynamics are fixed in exponential phases of growth [1, 2]. While this framework has yielded a rich mathematical literature establishing general criteria for community stability and coexistence [3–8], it relies on environments that are effectively clamped. Such models are inconsistent with the dynamics of many natural microbial communities, which instead assemble, grow, and disperse repeatedly in environments that change over the course of each growth cycle [9–12].

In these communities, interactions shift as the community matures through endogenous changes in local context, including resource depletion, metabolite buildup, spatial self-organization, and physiological state transitions [9–20]. Numerous studies have teased apart metabolic cascades [10], cross-detoxification and crossfeeding mechanisms [15, 17, 19–21], and chemotactic [14, 16] and diauxic [18] tradeoffs. These detailed mechanistic models can be used to build quantitative criteria for coexistence and stability, as in the case of static environments, but their system-specific nature makes it difficult to expand these criteria across distinct modes and mechanisms of community maturation.

In this work, we seek a mechanism-agnostic predictor of robustness that generalizes across many distinct modes of community maturation. We study the stability of microbial communities undergoing repeated growth–dispersal cycles, as in serial dilution experiments or natural colonization of non-replenishing, nutrient-rich patches. Within each growth cycle, resource levels, metabolites, and spatial structure evolve, causing species interactions to change dynamically over time. We focus on robustness under stochastic dispersal, modeled as demographic noise in how each new environment is seeded by a finite number of individuals, leading to multiplicative fluctuations in species abundances from cycle to cycle.

We introduce *co-growth*, the temporal overlap in rapid growth of different species during a growth cycle, as a mechanism-agnostic quantity that predicts community robustness. If species grow at similar rates in time during a growth cycle they have high levels of co-growth; if species grow at differential rates in time during a growth cycle they have low levels of co-growth (Fig. 1). The resulting temporal separation of growth resembles classical temporal niche partitioning [22, 23], but with a key distinction: here the temporal structure arises endogenously within each growth cycle due to temporally varying interactions between microbes rather than being imposed externally by environmental variation. Because co-growth is defined in terms of total growth rates (e.g., summed over all spatial locations or physiological states), it can be measured and compared across mechanistically distinct systems without requiring detailed knowledge of the underlying interactions.

**FIG. 1.**
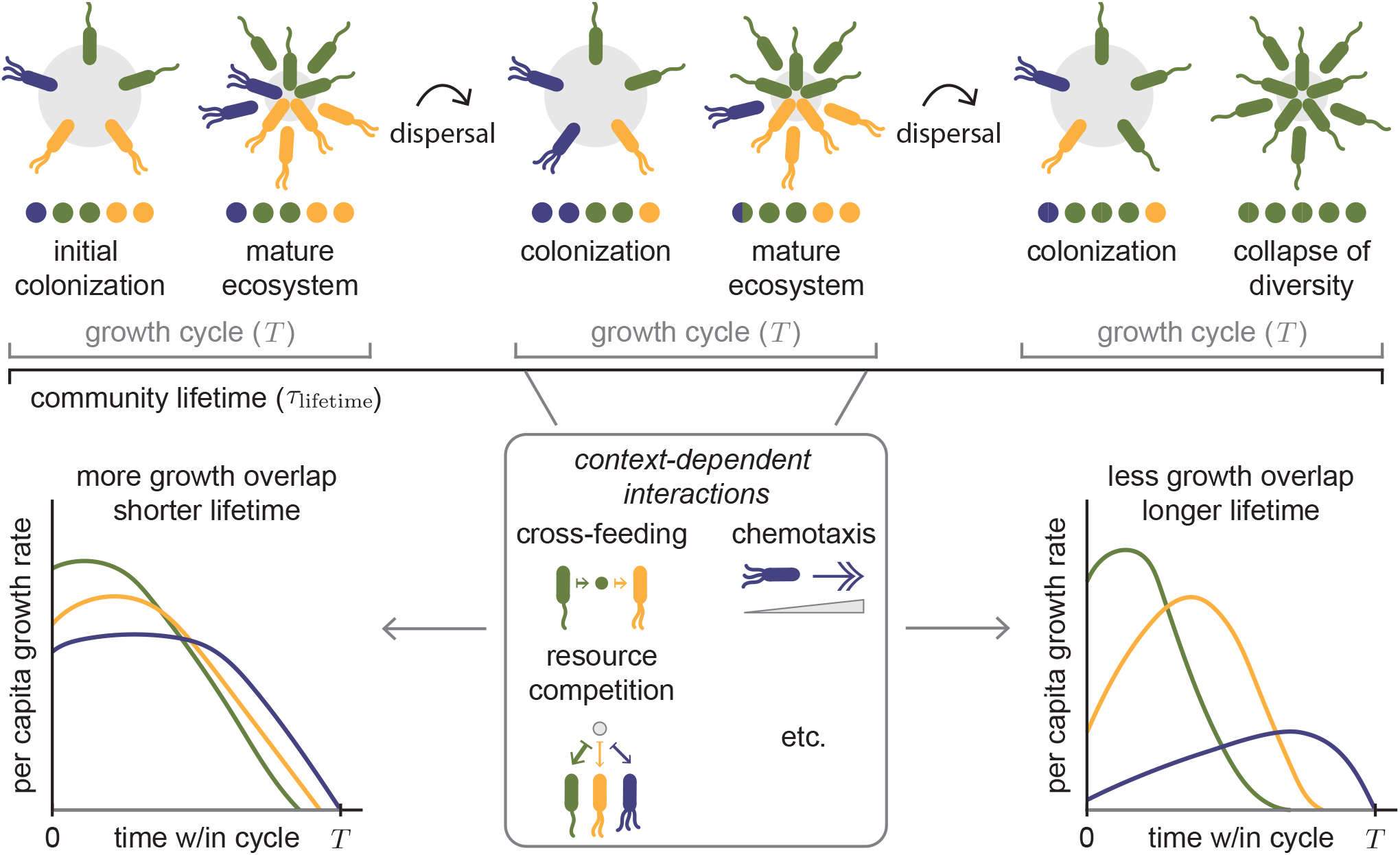
Minimizing co-growth increases community robustness in fluctuating environments. We model communities that coexist stably and undergo repeated growth–dispersal cycles. Within each cycle, species transition through context-dependent growth phases shaped by spatial structure, metabolite exchange, and other ecological interactions. Demographic noise during reseeding introduces small compositional shifts that can accumulate toward extinction. The resulting community lifetime (𝒯_lifetime_) reflects robustness to such drift.

We establish this predictive relationship using both mechanistic models and a phenomenological many-species framework, demonstrating that co-growth determines the scaling behavior of average community lifetime from within-cycle growth trajectories alone. We further show that this relationship follows from a general theoretical argument: the dynamics of community composition across cycles map onto a noise-driven escape problem, in which the mean time to extinction depends exponentially on a barrier height set by co-growth. This Arrhenius-like scaling does not rely on the specific mechanism generating temporal separation, suggesting that the predictive power of co-growth extends beyond the particular systems analyzed here.

## I. A GROWTH-DISPERSAL MODEL FOR COMMUNITY DYNAMICS

We consider communities that coexist stably while undergoing repeated growth-dispersal cycles in the wild or in serial dilution experiments in the lab. Within each growth cycle, a community colonizes a resource-rich medium (e.g., a chitin particle or a fresh plate) and grows in biomass. Species grow and interact through context-dependent processes such as the formation of spatial structure, the buildup or depletion of metabolites (nutrients or toxins), and other mechanisms discussed in the literature (Fig. 1) [10, 14–20]. These interactions can push each species through different growth phases over the course of a single cycle.

There are two natural timescales in this setting. Within one growth cycle, species densities and metabolites deterministically change until growth effectively stops at time *T*. We describe the density of species *α*, 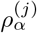, and metabolite *s*,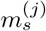, in cycle *j* by,

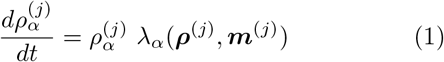

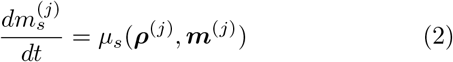

where *λ*_*α*_ (…) is the per-capita growth rate and *µ*_*s*_(…) denotes the mechanism-driven depletion or creation of metabolites, capturing how context-dependent interactions shape the dynamics of species and metabolites (Fig. 1). Note that given the assumption of deterministic dynamics and the initial conditions for metabolites in each cycle remaining the same, the population density of the species at the end of a cycle depends solely on their initial values, 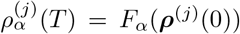, which defines the discrete map *F*_*α*_ (…) for a species *α*. In other words, the mechanistic details of growth within a cycle are encapsulated in this coarse-grained mapping, dependent only on the initial community composition.

Across many such cycles, the overall community composition slowly drifts in response to dispersal. At the end of each growth cycle, a small fraction, *δ*, of the community disperses and colonizes a new patch

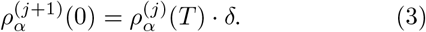

We assume that the community has a steady state ***ρ***^*^, such that 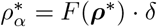 for all *α*.

With this separation of timescales, we treat within-cycle growth as effectively instantaneous and deterministic, relative to the gradual accumulation of stochastic effects across cycles. We model stochasticity in dispersal as demographic noise in how the seed community is assembled: each new environment is seeded by a finite number of individuals, so the realized species fractions fluctuate around their deterministic values with noise that scales with abundance. These sampling effects act as multiplicative perturbations to abundances from cycle to cycle. The product of many such factors is well approximated by a log-normal term, motivating the following form of stochasticity:

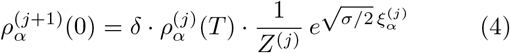

where *σ* is the magnitude of noise, 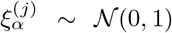, and 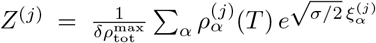 is a normalization factor ensuring that the total community biomass,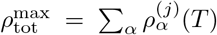, remains the same after the noise addition. The demographic noise pushes away the community from its steady state composition, and on long timescales can lead to the extinction of species. As a result, long-term community persistence becomes a problem of resisting noise-driven escape from the coexistence fixed point set by the deterministic within-cycle dynamics.

This motivates a natural notion of community robustness at the cross-cycle timescale. We characterize robustness by the community lifetime, 𝒯_lifetime_, the number of cycles before demographic noise during dispersal drives at least one species to extinction. A species *α* is considered extinct once its abundance after dilution, *ρ*_*α*_(*T*) · *δ*, falls below a threshold density *ρ*_extinct_ (with 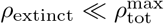), which we will call the extinction boundary. From the perspective of demographic noise driving an escape from coexistence, the average community lifetime (over noise realizations at fixed initial conditions and parameters) can be interpreted as the characteristic time required for the system to reach and cross the extinction boundary.

Intuitively, this long-timescale robustness must link back to the interaction dynamics operating within a single growth cycle. We establish this link through cogrowth, that is, the extent to which species grow at the same time over a cycle. We define a co-growth metric as a divergence measure (similar to the Kullback–Leibler divergence) for the per capita growth rates of different species as a function of time, *λ*_*α*_ (*t*). We require this measure to satisfy the following properties: (i) the measure is minimal when the growth profiles are identical throughout the cycle; (ii) the measure has a very large value when each species has a period during which it can grow while the other species stops growing (or has an almost zero growth rate); (iii) we assign greater weight to times when the absolute growth rates of both species are higher, and assign less weight when both species have almost no growth; and (iv) the measure should be dimensionless and symmetric under a permutation of species.

While several metrics might satisfy these criteria we use the time average (with a weight) over a cycle of the ratio between growth rates. With two species, this is simply:

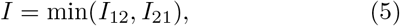

Where

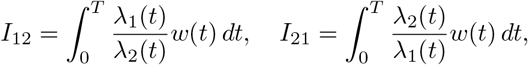

and *w*(*t*) is the weight applied to the average. Here, we choose the growth rate of the total biomass as the weight,

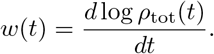

For further discussion about the *N* > 2 species case, see Sec. V and SI Sec. S3.

This co-growth metric *I* attains its minimum value when *λ*_1_(*t*) ∼ *λ*_2_(*t*) over the whole cycle. In contrast, it becomes very large when there is perfect temporal separation, such that *λ*_1_(*t*) > 0 and *λ*_2_(*t*) ≈ 0 at some time *t*, and *λ*_2_(*t*^′^) > 0 and *λ*_1_(*t*^′^) ≈ 0 at another time *t*^′^. It is worth noting that, by the change of variables *d* log *ρ*_tot_ = (*dρ*_tot_*/ρ*_tot_), this measure is equivalent to the invasibility metric put forward in [24].

In spatially heterogeneous settings, the same definition applies after replacing *ρ*_*α*_ (*t*) by the total biomass integrated over space

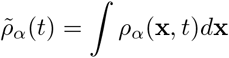

and *λ*_*α*_ (*t*) by the corresponding biomass-weighted effective per-capita growth rate

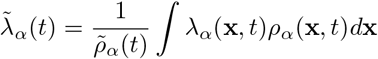

(see SI Sec. S1.B for an explicit example).

A key feature of this approach is that this metric of co-growth coarse-grains over all mechanistic complexity, integrating over biomass across space and physiological states. This coarse-graining is not a loss of information but enables generalization. As we will demonstrate, cogrowth, as such, connects to average community lifetime independent of underlying mechanism.

## II. SPATIAL PATTERNING AS A MECHANISM TO STRUCTURE GROWTH IN TIME

To understand how co-growth can provide a principle for the robustness of communities assembled through complex, not inherently temporal, mechanisms, we examine a case study of interactions mediated through complicated spatial structuring driven by chemotaxis.

Bacterial chemotaxis couples multiple cellular processes, including growth, swimming, nutrient uptake, and the sensing of chemical gradients [25]. Each of these processes depends on the cell’s physiological state and local environment, resulting in nonlinear dynamics at a population level. Movement and growth in the community are governed by distinct molecular species: a primary nutrient (e.g., a carbon source) fuels cell growth, while a chemoattractant (e.g., an amino acid present at much lower concentration) is sensed by cells and provides directional cues for migration [25]. Although these can sometimes be the same molecule, they need not be, and the interplay between independent nutrient and attractant fields, together with diffusion and cell motility, produces rich spatiotemporal dynamics in expanding colonies [14, 25].

Many studies have shown that microbial species can coexist through complex spatial structuring in which nutrient diffusion, growth, and chemotaxis interact to create gradients that sustain an ecosystem [14, 16, 26–30]. As a concrete example of such mechanisms, we focus on a system in which coexistence arises from a complex balance between motility and growth (Fig. 2a) as seen experimentally [14, 16]. When two species are inoculated at the center of a circular plate with a uniformly distributed nutrient and chemoattractant, the species can spontaneously segregate into proximal and distal territories (Fig. 2b). While a trade-off between growth rate and migration speed is a necessary qualitative condition for this coexistence, the precise criterion is far more complex: the rates of nutrient depletion and attractant consumption, their respective diffusion coefficients, and the nonlinear coupling between local growth and chemotactic drift all jointly determine whether and how species coexist.

**FIG. 2.**
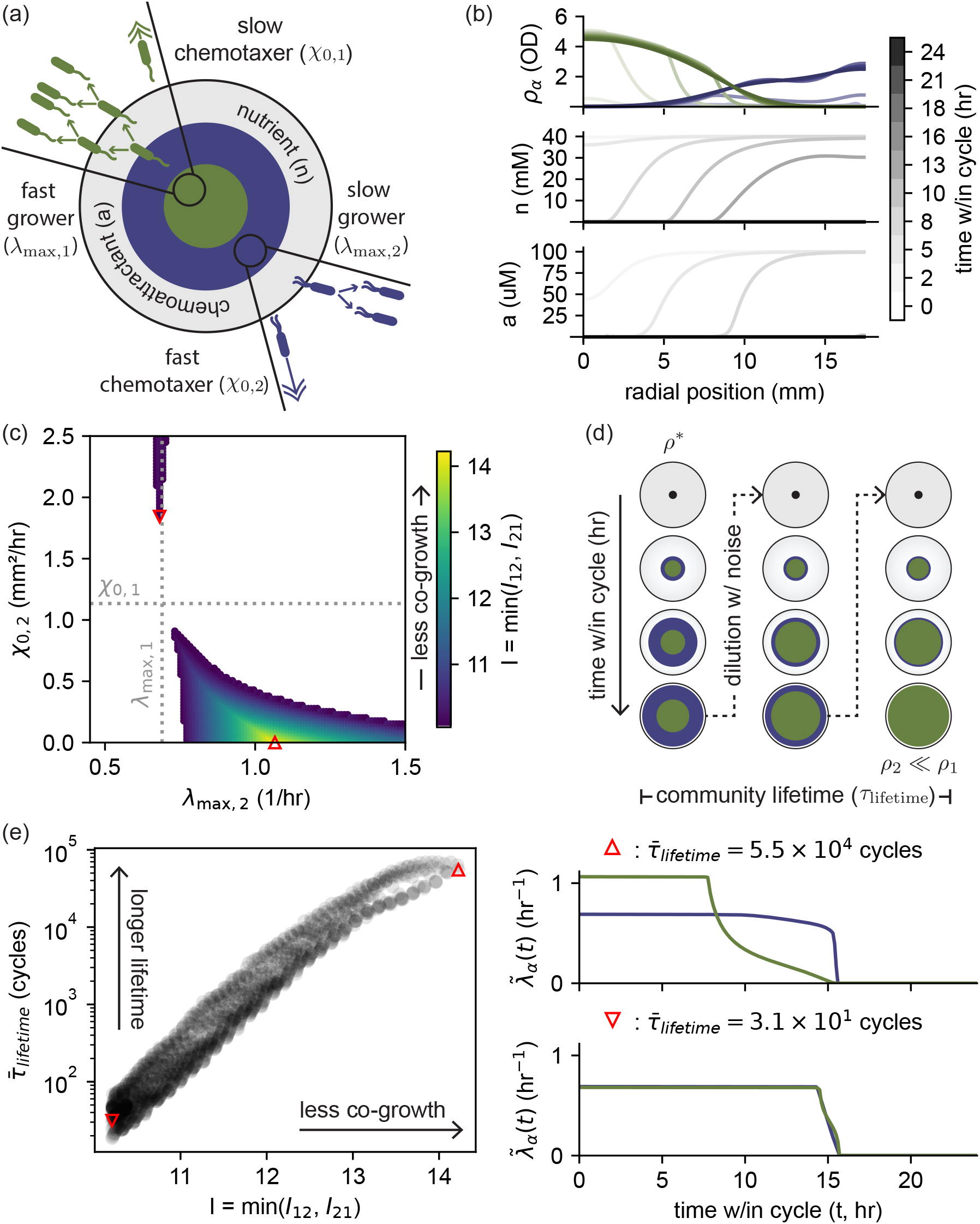
Complex spatial patterning can be viewed as a mechanism to reduce temporal overlap in growth periods. **(a)** Spatial dynamics inspired by the experimental framework of [16]. Each species is characterized by two traits that govern its interaction with chemoattractant and nutrient fields shared by both species: their maximal growth rate *λ*_max,*α*_ and their chemotactic coefficient *χ*_0,*α*_. **(b)** Spatial density profiles of the two species (green, blue) plotted over time. Darker shades denote later times. **(c)** Coexistence landscape across many “strategies” defined by the trait set {*λ*_max,*α*_, *χ*_0,*α α*=1,2_}. Parameters for species 1 are fixed (dotted lines), while species-2 parameters were evenly sampled on a grid across the full range: *λ*_max,2_ ∈ [0.5, 1.5] and *χ*_0,2_ ∈ (0, 2.5]. Only strategies yielding coexistence are plotted. Heatmap indicates the degree of co-growth for each strategy. **(d)** Community lifetime is calculated by running the experiment *in silico* with noisy dilution starting at the steady state, ***ρ***^*\**^, until one of the two species goes extinct, *ρ*_2_ ≪ *ρ*_1_. **(e)** (left) The average community lifetime 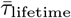, averaged over noise realizations at fixed initial conditions and parameters, as a function of co-growth. Strategies that promote stronger temporal separation (lower co-growth) yield longer-lived communities. (right) Example global per-capita growth rate,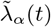, trajectories for a long-lived and a short-lived strategy with initial abundance at the fixed point, ***ρ***^(0)^ = ***ρ***^*\**^, illustrating temporal separation of growth within the growth cycle.

Following the growth-expansion framework of [25], we model this two-species spatial community in radial geometry by coupling the density of each species, *ρ*_*α*_ (*r, t*) (*α* = 1, 2), to a nutrient field, *n*(*r, t*), and a chemoattractant field, *a*(*r, t*). Growth is driven by nutrient availability, while directed movement is guided by the attractant gradient. Because the nutrient and attractant have different concentrations and distinct diffusion and consumption rates, they develop distinct spatial profiles, and their combined influence shapes the population dynamics (for further discussion of this model, see SI Sec. S1).

Each species is characterized by a maximal growth rate *λ*_max,*α*_ and a chemotactic coefficient *χ*_0,*α*_. The dynamics are:

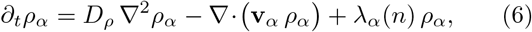

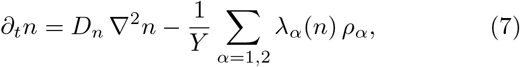

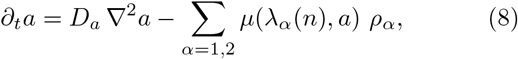

where the (Monod) local growth rate is

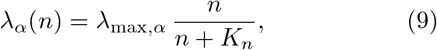

*Y* is the growth yield, and *D*_*ρ*_, *D*_*n*_, and *D*_*a*_ are diffusion coefficients for cells, nutrient, and chemoattractant, respectively [25]. Directed motion along chemoattractant gradients enters via the drift velocity

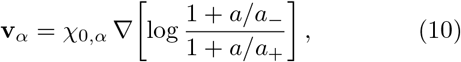

where *a*_−_ and *a*_+_ (the lower/upper Weber offsets) set the receptor sensitivity range [25]. Chemoattractant consumption is modeled as an uptake function that increases with growth rate and saturates with chemoattractant concentration,

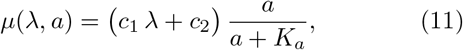

with experimentally fit constants *c*_1_ and *c*_2_ and Monod constant *K*_*a*_ [25].

The coupled dynamics generate the annular spatial pattern observed experimentally (Fig. 2a,b). Initially, both species compete locally for the shared nutrient near the inoculation site, and the faster grower (green) dominates there. Nutrient and chemoattractant consumption creates a depleted zone around the center. The species with higher chemotactic sensitivity (blue) follows this self-generated attractant gradient outward, establishing a dense ring at the colony periphery and restricting nutrient supply to the inner population. After the nutrients on the plate are depleted, marking the end of one cycle, the species disperse to another plate inheriting the ratio of abundance as summed across the full plate.

We quantitatively investigated the conditions for coexistence by parameterizing each community *strategy* by the maximal growth rates and chemotactic coefficients of the two species. We swept a two-dimensional parameter space over the species 2 traits, (*λ*_max,2_, *χ*_0,2_), while holding the species 1 traits (*λ*_max,1_, *χ*_0,1_) fixed, along with all other parameters (i.e., diffusion coefficients, nutrient/chemoattractant kinetics, and the chemotactic coupling to gradients). Coexistence appears primarily in specific regions where no species is superior in both traits (Fig. 2c): in the top-left quadrant, species 1 is the faster grower (green, *λ*_max,1_ > *λ*_max,2_) and species 2 is the better chemotaxer (blue, *χ*_0,2_ > *χ*_0,1_), while in the bottomright quadrant the roles are reversed (*λ*_max,2_ > *λ*_max,1_, *χ*_0,1_ > *χ*_0,2_). (We also observe a narrow sliver of coexistence even when species 2 exceeds species 1 in both traits, *λ*_max,2_ > *λ*_max,1_ and *χ*_0,2_ > *χ*_0,1_, suggesting that additional mechanistic features of the full model can weakly counterbalance a naive “dominance in both traits” expectation; we leave mechanistic interpretation of this regime to future work and focus on the phenomenological prediction based on co-growth here.) The shape of these coexistence regions is set sensitively by all the other parameters; small changes in physiological or physical parameters can move the system into or out of the coexistence region in ways that are not easy to predict from the full mechanistic model alone.

We therefore ask whether the principle of minimizing co-growth provides a simpler way to organize these outcomes. In this view, the complex spatial patterning described above is merely a mechanism to reduce temporal overlap in rapid growth. Quantifying co-growth across strategy space shows that as strategies move toward the edges of the coexistence region, the two species tend to grow at similar times and co-growth increases (darker colors). As strategies move toward the interior, growth in space and time separates more cleanly and co-growth decreases (lighter colors) (Fig. 2c).

We next ask how robust these coexisting communities are, beyond the binary question of whether they coexist. To do so, we simulate growth and dispersal cycles to find the stable composition (***ρ***^*^) of each coexisting community, then introduce fluctuations with fixed fluctuation intensity at each dispersal and measure community lifetime (𝒯_lifetime_), i.e. how many cycles elapse before one of the two species goes extinct (Fig. 2d). Applying this lifetime computation across all of parameter space reveals an approximately log-linear relationship between our cogrowth metric and average community lifetime, averaged over noise realizations at fixed initial conditions and parameters, (Fig. 2e). Strategies that yield long lifetimes use spatial structure to strongly separate growth in time: the green population blooms locally near the center and remains confined there, while the blue population must first swim outward before it can grow strongly, so their periods of rapid growth barely overlap. In strategies with shorter lifetimes, although coexistence is still possible, the two species tend to grow at overlapping times, cogrowth is higher, and extinction occurs sooner (Fig. 2e).

## III. TOXICITY AS A MODULATOR OF TEMPORAL GROWTH STRUCTURE

While interactions through chemotaxis provide a compelling demonstration of how spatial structure can be seen as in service of minimizing co-growth, we seek to show that this principle applies equally well to drastically different mechanisms of community maturation. As a second example, we examine a cross-feeding model based on the first step of denitrification, commonly done by microbial communities in soils [17, 20]. In this system, two generalist strains can metabolize nitrate 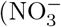, denoted *n*_*A*_(*t*)) into nitrite 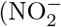, denoted *n*_*I*_ (*t*)) and nitrite into nitrous oxide (NO).

Within soils across the globe, there exists a trade-off between two genotypes of denitrifiers as soil pH shifts: *nar* gene abundances are more present as pH decreases while *nap* gene abundances are more present as pH increases [20]. Crocker *et al*. further showed that, despite the global trend of Nar^+^ strains with low-pH soils, Nar^+^ strains grow poorly in low-pH monoculture and instead require a Nap^+^ strain for enrichment [20]. In their experiments, low pH makes nitrite toxic to Nar^+^ strains, and the accompanying Nap^+^ strain is required to prevent nitrite accumulation by rapidly consuming nitrite [20]. Crocker *et al*. propose that the interactions balance at low pH because the Nar^+^ strain outcompetes in the consumption of nitrate but is a poor nitrite consumer, leading to to nitrite buildup and self-toxification; the Nap^+^ strain then helps the community recover by strongly consuming nitrite [20].

From a classic consumer-resource model perspective, the dominance of the two species on different metabolites should be sufficient for coexistence, yet toxicity dynamics, such as those described above, are not uncommon [19, 31]. Previous work has examined the ways that cross-detoxification interactions can strengthen mutualism in a cross-feeding setting [32], however, the role of toxicity more broadly in community maturation and robustness remains unclear.

We use a consumer resource model similar to [17], but add in a graded toxicity effect of nitrite on one of the species inspired by the toxicity effects seen on Nar^+^ strains (hereafter referred to as species 1, *ρ*_1_(*t*)) as pH is decreased while Nap^+^ strains (hereafter referred to as species 2, *ρ*_2_(*t*)) are unaffected [20] (Fig. 3a).

**FIG. 3.**
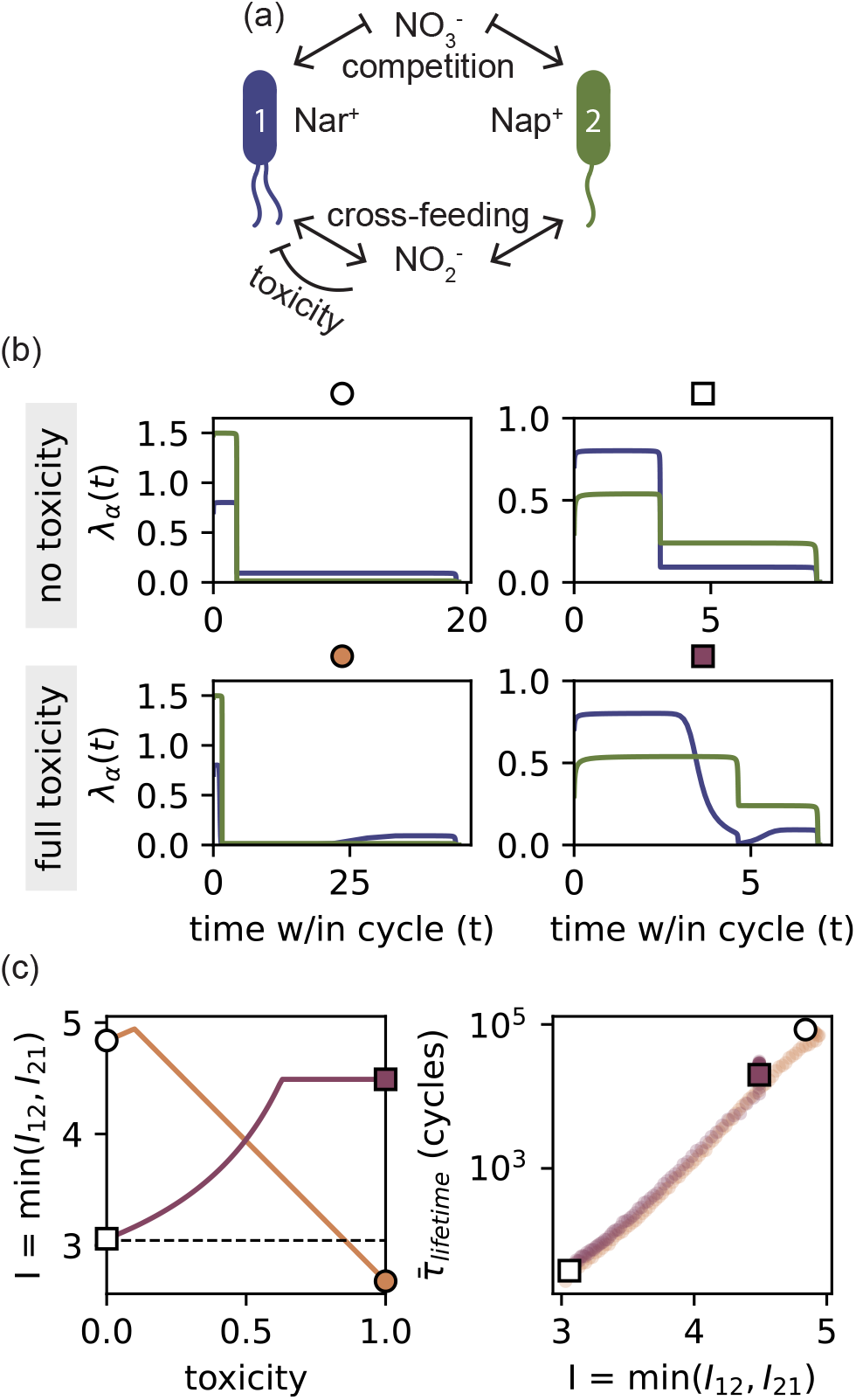
Toxicity modulates temporal growth structure in a cross-feeding denitrification model. **(a)** Schematic of the model: two generalist species consume nitrate (*n*_*A*_) and nitrite (*n*_*I*_), with species 1 (Nar^+^) experiencing a graded toxic effect from nitrite [20]. The toxicity parameter ranges from 0 (no effect) to 1 (maximal effect). **(b)** Per-capita growth rate trajectories with initial abundance at the fixed point, ***ρ***^(0)^ = ***ρ***^*\**^, for the two species (blue, species 1; green, species 2) plotted as a function of time, shown for two communities (tan circle and burgundy triangle) under no toxicity (toxicity = 0) and full toxicity (toxicity = 1). The two communities represent opposite toxicity-co-growth trends: in one (tan circle), toxicity increasingly suppresses species 1 to a point where species 2 outcompetes it; in the other (burgundy triangle), toxicity amplifies the temporal separation between the two species’ growth phases, reducing co-growth. **(c)** Co-growth, measured as *I* = min(*I*_12_, *I*_21_), plotted as a function of toxicity (left), and average community lifetime 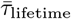, averaged over noise realizations at fixed initial conditions and parameters, as a function of co-growth. (right). The dashed line in the left plot marks the coexistence threshold, communities that fall below it lose stable coexistence, have no well-defined community lifetime, and therefore are not plotted on the right. Co-growth reliably tracks robustness in both cases: more cogrowth corresponds to shorter average community lifetimes, less co-growth to longer ones.

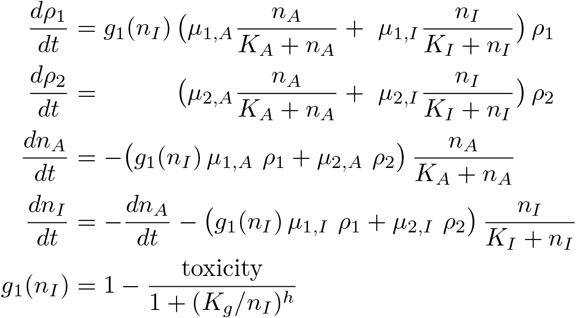

The strategy of a community is defined by the resource-specific maximum growth rates, {*µ*_*α,A*_, *µ*_*α,I*_}_*α*=1,2_, while the half-saturation constants (*K*_*A*_, *K*_*I*_, *K*_*g*_) and the Hill coefficient for the gain function (*h*) are held constant. Toxicity is introduced as a graded parameter between 0 (no toxicity) and 1 (maximal toxicity) (for further discussion of this model, see SI Sec. S2).

The effect of toxicity on co-growth is not uniform: it depends on the competitive landscape set by the background growth rates. To show that co-growth remains a reliable predictor of robustness regardless of where in the interaction landscape one is, we examine two characteristic communities with distinct growth rate parameters {*µ*_*α,A*_, *µ*_*α,I*_} _*α*=1,2_ (Fig. 3b,c).

In one community (tan circle; Fig. 3b,c), increasing toxicity progressively suppresses species 1’s growth until coexistence breaks down entirely. In the other (burgundy triangle; Fig. 3b,c), toxicity has the opposite effect: it constrains species 1’s growth during a middle period when nitrite has accumulated, giving species 2 a stronger growth advantage and causing a temporal restructuring that manifests as reduced co-growth and longer average community lifetime (Fig. 3c). In both cases, co-growth remains a reliable predictor of robustness (see SI Sec. S2.F for a broader exploration of parameter space).

This example reveals a key advantage of the co-growth framework: it enables characterization of community dynamics without requiring complete mechanistic knowledge. Here, the underlying model involves a complex interplay of nutrient competition, cross-feeding, and toxicity (which also implies detoxification through consumption). Depending on where you are in parameter space, small changes to one growth rate or the toxicity level can have very different effects on the robustness and stability of the community. Even without a full dynamical model, one could assess the local gradient of change in robustness given the metabolic background consumption rates of the species, by buffering the toxic compound and measuring how growth curves change. The co-growth metric calculated from these growth curves alone would reveal whether toxicity stabilizes, destabilizes, or has no effect on the community, independent of the detailed mechanisms at play.

## IV. LINKING CO-GROWTH TO LIFETIME THROUGH ARRHENIUS’S LAW

To explain the surprising log-linear relationship between co-growth and lifetime, it is useful to think of the dynamics of the community composition as described by the log ratio, 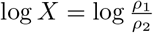.

With demographic noise, the community composition performs a random walk with a restoring drift in the space of log abundance ratios. The dynamics of the log ratio across cycles can be written as,

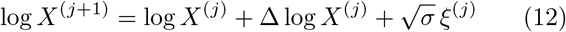

where Δ log *X*^(*j*)^ captures the deterministic dynamics, or the net effect of interaction<u>s</u> within one growth cycle in the absence of noise, and 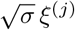 captures the stochastic component. Note that since the log-ratio captures the single compositional degree of freedom in a two-species community, the two independent per-species noise terms described in Eqn. 4 reduce to a single effective scalar noise *ξ*^(*j*)^ ∼ 𝒩 (0, 1) acting on log *X*; more generally, an *N*-species community has *N* − 1 compositional degrees of freedom and thus *N* − 1 independent noise terms. In this view, interactions within a cycle set Δ log *X*^(*j*)^, and therefore determine how we measure co-growth, while the long-term community composition, and its noise-driven departures from coexistence, play out over many cycles.

Recast, the dynamics of the log ratio can be thought of as a gradient flow on an effective potential,

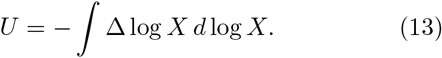

Intuitively, *U* is a landscape in composition space: the coexistence state sits at a local minimum of *U*, and the deterministic dynamics move the community composition downhill toward this minimum (Fig. 4). From a ratio perspective, the extinction boundary can now be described by an extinction ratio, *X*_extinct_. Species 1 is considered extinct when the compositional fraction, 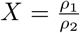 is greater than a specified threshold, *X*_extinct_ ≫ 1. Symmetrically, Species 2 is considered extinct when: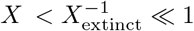.

**FIG. 4.**
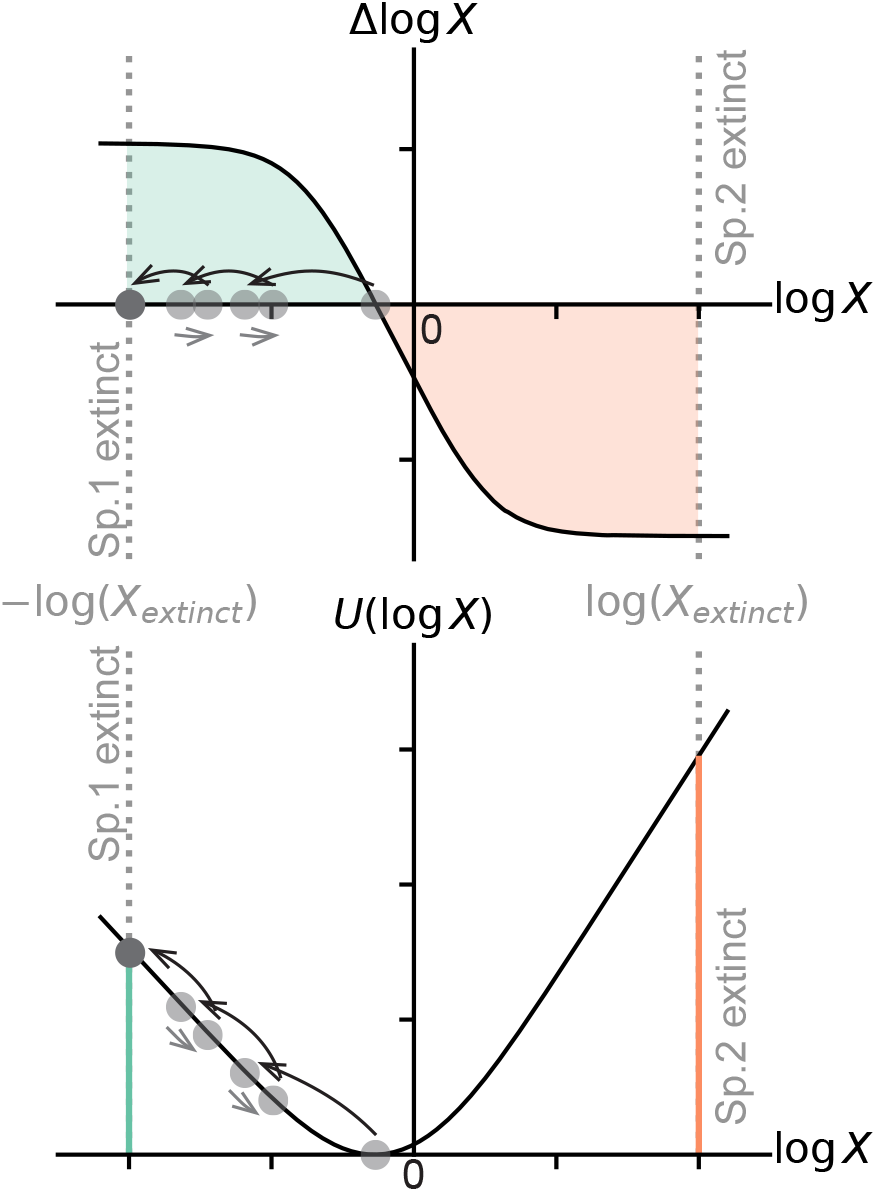
Average community lifetime can be linked to a metric of co-growth through an Arrhenius approximation. **(top)** The change in community composition (Δ log *X, X* = *ρ*_1_*/ρ*_2_) over one growth cycle as a function of the initial composition of the system (log *X*) in the absence of noise describes the deterministic dynamics of the system over multiple cycles. Dotted lines mark the extinction boundary (± log *X*_extinct_). **(bottom)** Energy *U* is defined as the integral of the change in composition over the course of one cycle, *U* = − Δ log *X d* log *X*.

At first sight, one might expect the time needed to escape from the coexistence state and reach an extinction boundary due to noise to depend on all the detailed features of this landscape *U* and therefore have more to do with the path taken across cycles than any property of the dynamics within one cycle (like co-growth). However, classic results related mathematically to work in thermodynamics by Arrhenius can be used to show that this is not the case when the coexistence state is sufficiently stable and noise is weak [33]. In that regime, the noisy dynamics can be approximated by a saddle point evaluation of the path integral over escape trajectories, so that the mean escape time depends primarily on the height of the lowest barrier separating the stable state from the nearest saddle point or absorbing boundary (in this case species extinction). The detailed shape of the trapping potential near the minimum only affects prefactors, while the dominant exponential dependence of the average community lifetime is set by the barrier height [34, Ch. XIII, §2]. Therefore, the log of the averaged community lifetime 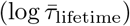 is represented by

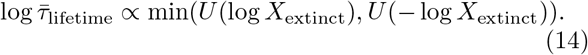

The boundary energy, *U* (±log *X*_extinct_), ultimately ties average community lifetime back to the timescale of dynamics within one growth cycle because it integrates over the detailed interactions to provide an overview. Pushing the extinction boundaries to log *X*_extinct_→∞, the extinction energy can be approximated as proportional to the invasibility of the endangered species.

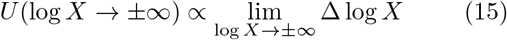

Previous work [24] has shown that the invasibility simplifies to:

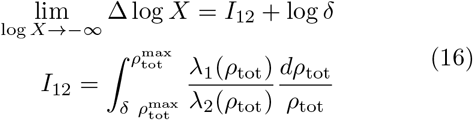

returning back to the quantitative measure of co-growth we had motivated before (Eqn. 5). This adds the final link to explain the approximate log-linear relationship between average community lifetime and co-growth via Eqn. 14.

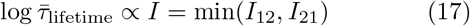

Taking the minimum of the two invasibility metrics is now explained as corresponding to selecting the lower of the energy barriers at the extinction boundaries.

## V. A PHENOMENOLOGICAL MODEL OF MANY SPECIES

In the previous sections we have demonstrated the generality of co-growth as an indicator of community robustness across multiple specific interaction mechanisms. In all of these cases, the unifying principle is that the growth curves within a growth cycle alone, through the co-growth metric, are sufficient to characterize both coexistence and robustness, without requiring knowledge of the complex underlying biological mechanisms. While the mechanistic examples shown previously are for *N* = 2 species, this framework generalizes to arbitrary *N* > 2 species systems. For the full theoretical framework extending co-growth as a predictor to *N* > 2 species, see SI Sec. S3.

In brief, we extend the co-growth metric to *N* > 2 species by defining *I*_*α*_ as the invasion growth of species *α* at low abundance into an arbitrary (*N* − 1)-species subcommunity. Specifically, let 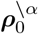 denote any initial condition with 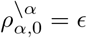 and 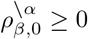 for all *β* ≠ *α*, where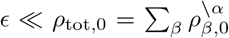. The invasion growth of species *α* into this subcommunity is

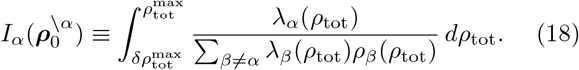

Analogous to the two-species case, where the co-growth metric (Eqn. 5) takes the minimum over both invasion directions, the community co-growth for *N* > 2 species is

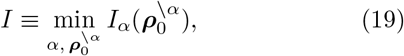

where the minimum now runs over all species *α* and all possible compositions 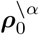 of the (*N* − 1)-species subcommunity. As before, this quantity sets the barrier height of an energy profile along the escape trajectory, directly linking co-growth to average community lifetime.

To demonstrate this, we constructed a phenomenological model in which growth rates are decoupled from a specific biological mechanisms. Each species follows the general rate equation 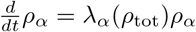 (Eqn. 1), where the per-capita growth rate *λ*_*α*_ is an arbitrary function of total community biomass *ρ*_tot_, following the communitystate reparameterization of time introduced in [24]. This growth function is discretized into *M* equal biomass bins with the width Δ*ρ*, with *λ*_*α*_ (*ρ*_tot_) = *G*_*α,b*_ in the *b*th bin (*b*Δ*ρ*≤ *ρ*_tot_ *<* (*b*+1)Δ*ρ*), so that the community strategy is fully encoded in an *N* × *M* growth rate matrix *G* whose rows index species and whose columns index successive biomass bins. For small Δ*ρ*, the species abundances are well approximated by the discrete update rule

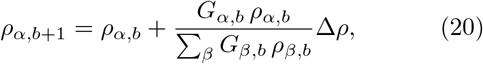

where *ρ*_*α,b*_ is the abundance of species *α* in bin *b* and Δ*ρ* is the community biomass increment per bin (for further discussion of this model, see SI Sec. S4).

Using this model, we built an ensemble of five-species communities (each defined by a 5 × 5, *G* matrix) via replica-exchange Monte Carlo, retaining only community with full coexistence and sampling broadly across cogrowth values (see SI Sec. S4.D). Rare-event sampling ensured adequate coverage of communities with higher co-growth values. Across this ensemble, we recapitulate the co-growth - robustness relationship observed in twospecies mechanistic systems (Fig. 5): communities with lower co-growth are systematically more robust to demographic fluctuations.

**FIG. 5.**
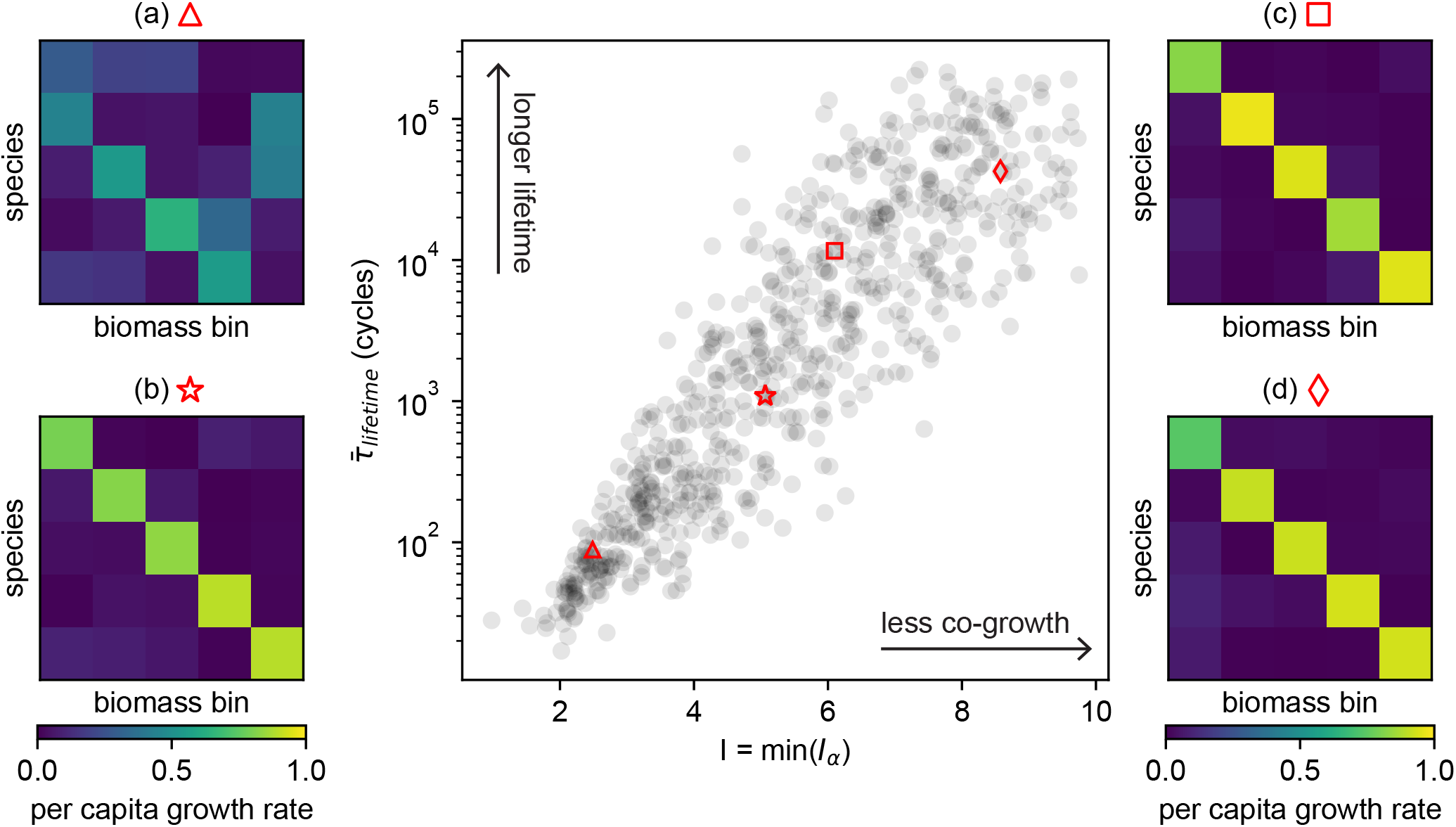
The co-growth–robustness relationship generalizes to communities with *N* > 2 species. **(center)** Average community lifetime 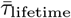, averaged over noise realizations at fixed initial conditions and parameters, as a function of cogrowth for an ensemble of five-species communities generated from random 5 × 5 growth matrices *G*, where each entry *G*_α,*b*_ specifies the per-capita growth rate of species α in biomass bin *b*. Using a phenomenological model of context-dependent interactions, we sample matrices and retain only those that support full coexistence of all five species. Average community lifetime increases systematically as co-growth decreases (horizontal axis: *I* = min(*I*_α_)), revealing the same robustness principle found in two-species systems. **(a–d)** Example growth matrices from strategies with different levels of co-growth, as indicated by the corresponding markers on the central plot. As co-growth decreases, species increasingly partition their growth across biomass bins, illustrating stronger temporal separation of growth.

Minimizing co-growth, however, presents a trade-off. The temporal separation of growth in time coincides with species specializing along a dynamic cascade and therefore removing any single species forces the remaining species to compensate for a missing link in the temporal sequence, slowing down the overall growth cycle (Fig. 6), a dynamic we call “arrested development”. For instance, the loss of a species that has specialized to complete a particular step in a cascading biochemical process may then slow down the overall community development because a less skilled species has to take on that role to compensate. Crucially, this slow down is most severe in the low co-growth communities that are otherwise most robust (i.e. longer average community lifetime), revealing a fundamental tension between robustness to demographic fluctuations and resilience to species loss. The effect of arrested development reveals the fundamentally cooperative nature of these ecosystems: each species acts as a temporal keystone species, preparing the ecological context for species that follow, and is irreplaceable even if other species nominally could perform similar functions.

**FIG. 6.**
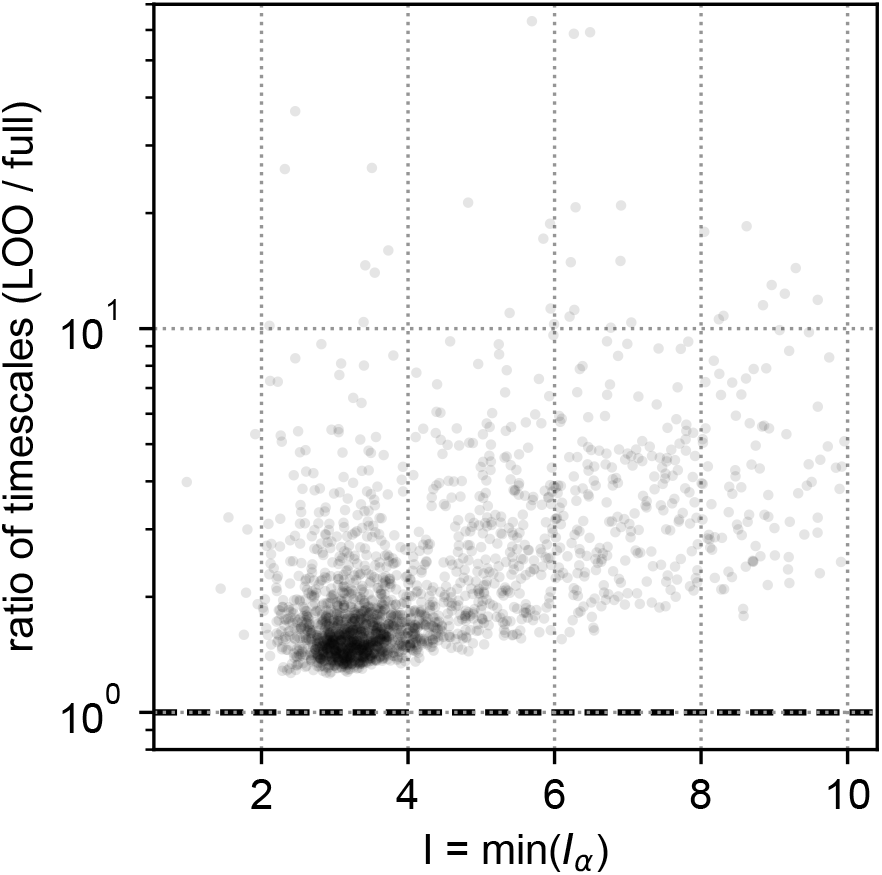
Removing any single species slows the community’s progression through a growth cycle, with stronger effects in high co-growth communities. For each five-species community in the ensemble introduced in Fig. 5, we compute the time to traverse all biomass bins twice: once using the full community at its steady-state composition, and once using the same steady-state composition but with a single species removed (“leave-one-out,” LOO). For each community we evaluate the LOO cycle time for all five possible single-species deletions and record the maximal slow-down, 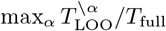. Cycle times are calculated from the per-capita growth rates in each biomass bin *b* as 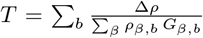, so that removing a species reduces the biomass contributing to growth in each bin. The plotted ratio (LOO / full) is therefore always ≥1. Communities with higher co-growth exhibit larger maximal increases in timescale upon species removal.

## VI. DISCUSSION

### 1. Conclusions

This work shows that minimizing co-growth provides a unifying principle for predicting and comparing the robustness of vastly different ecological systems. Diverse mechanisms, such as spatial patterning, toxin or metabolite buildup, metabolic cascades, and physiological state transitions, can all be understood as routes to the same functional outcome, temporally separating rapid growth, so that different species peak at different times within the cycle. More broadly, this helps bridge a gap in microbial ecology; we have general mathematical criteria for community stability in clamped environments (i.e., where growth conditions and interactions are artificially fixed such as in a chemostat), but lack comparable principles for communities in growth-dispersal cycles, where resources, metabolites, and spatial structure evolve within each cycle and interactions shift as a result.

Our framework connects to the broader coexistence and niche-overlap literature [35–38], but addresses a distinct question: not simply whether species *can* coexist together, but *how robustly* they coexist under repeated growth-dispersal cycles with stochastic seeding. Unlike classical temporal niche partitioning, where environmental variation is imposed externally (e.g., seasonal fluctuations, diurnal cycles) to carve out distinct time windows for different species [22, 23], the temporal separation that emerges in low co-growth communities is generated from within: through spatial self-organization, metabolite buildup, and physiological state changes. In other words, the community creates its own temporal niches. Co-growth captures the net effect of these diverse processes on a single axis, making it possible to compare systems whose microscopic mechanisms are otherwise difficult to relate. This shift from externally imposed to internally generated temporal structure suggests that community robustness can be tuned by manipulating how species interact within a cycle rather than requiring carefully orchestrated environmental forcing.

### 2. Limitations of the current framework

An implicit assumption of our framework is that species interact through overlapping ecological niches, such that they compete for shared resources, metabolites, or space and their growth dynamics are entangled. The theory does not apply in regimes where species occupy effectively disjoint niches, for example, when each species consumes a distinct resource pool or grows in spatially separated patches, since coexistence in such cases is independent of co-growth. As such, this theory cannot be applied to situations such as obligate cross-feeding because although such species depend on one another for survival, their interaction effectively partitions resource use across complementary niches rather than introducing direct competition along a shared resource axis.

Another limitation lies in the fact that the current framework focuses on demographic noise during dispersal in serial-dilution-type systems. It does not yet fully address other sources of noise such as demographic or environmental fluctuations within growth cycles, or variability in the timing or strength of dispersal events (e.g. dispersal could stochastically occur before full nutrient depletion or the seeding biomass could fluctuate). In practice, we approximate the accumulation of these multiple sources of noise as a single log-normal perturbation applied at dispersal with the reasoning that the population experiences its strongest bottleneck, and thus its strongest stochasticity, at that moment. This approximation allows us to connect community robustness and co-growth through the Arrhenius approximation under weak, constant noise and stable coexistence. Incorporating this alternative sources of noise, including timevarying fluctuations, may alter the relationship between co-growth and robustness particularly near coexistence boundaries, however, we predict that the general trend of a separation of growth in time leading to longer average community lifetimes will remain unaffected.

### 3. Future directions beyond this work

The framework’s key utility is that it provides a mechanism-agnostic screening metric for robustness. Rather than building a separate stability argument for each mechanism, we ask whether a perturbation (e.g., toxin sensitivity, nutrient diffusion, motility) shifts the community toward more or less temporal overlap in rapid growth within a cycle. Computing co-growth does require within-cycle trajectories, measured experimentally when possible, or generated by a model, but the conceptual reduction is substantial: many mechanistic details matter primarily through their net effect on temporal separation. In the toxicity case, this lens clarifies why changing a single parameter can increase robustness by increasing temporal segregation of growth, even without complete knowledge of the metabolic network or spatial dynamics that mediate the effect.

Critically, co-growth can be inferred directly from growth curves through experimental manipulations such as varying spatial structure, buffering toxins, or modulating resource availability. This enables systematic exploration of parameter space to optimize community robustness without requiring detailed mechanistic models. Growth curves alone predict whether a manipulation will increase average community lifetime, making the framework experimentally tractable across diverse systems.

Another distinct expansion of this work would be to look at the evolution of communities in concert with dynamics that separate growth in time. Our study samples a variety of communities and evaluates the relationship between their robustness and the growth profiles of their constituent species, but it does not address the evolution of those communities. Some robust communities appear to require rare parameter configurations, suggesting that directed evolution or artificial selection might be necessary for their emergence in practice. This raises an intriguing question: when evolutionary dynamics or species immigration are introduced, can low co-growth communities arise spontaneously? In other words, what is the origin of highly robust communities in natural or synthetic settings? Further exploration into the evolution of these communities would benefit directly from the criteria established in this work because they offer a guiding principle for understanding the community-level robustness and individual-level phenotype.

## ACKNOWLEDGMENTS

We thank Martin Falk, Riccardo Ravasio, Mason Rouches, Darren Liu, Stefano Allesina, Akshit Goyal, Abby Skwara, and the Murugan group at large for helpful discussions. The authors acknowledge the University of Chicago’s Research Computing Center for computing resources, the National Science Foundation through the Center for Living Systems (PHY-2317138), and the NSF-Simons National Institute for Mathematics and Theory in Biology (National Science Foundation award DMS-2235451 and Simons Foundation award MPS-NITMB-00005320). M.C-W. acknowledges a Fannie and John Hertz Fellowship Award and the National Science Foundation Graduate Research Fellowship Program under grant number DGE 1746045. A.M. acknowledges support from National Institute of General Medical Sciences of the NIH under grant number R35GM151211 and the Schmidt Sciences Polymath award. A.V.N. acknowledges support from the Stanford Science Fellows program. T.H. acknowledges support from the National Institute of General Medical Sciences of the NIH under grant number R35GM152133. Any opinions, findings, conclusions, or recommendations expressed in this material are those of the authors and do not necessarily reflect the views of the National Science Foundation.

