## Supplementary Information for "Minimizing co-growth as a broad predictor of community robustness"

**CONTENTS**


S1. Spatial patterning as a mechanism to structure growth in time 4

S1.A. The model 4

S1.B. Calculating co-growth with spatial heterogeneity 6

S1.C. Community lifetime under demographic noise 8

S1.D. Numerical implementation and ensemble construction 8

S1.D.1. Spatial discretization and boundary conditions 9

S1.D.2. Time integration within one growth cycle 9

S1.D.3. Identification of coexistence parameter sets 10

S1.D.4. Constructing the $\Delta \log X$ map 10

S1.D.5. Co-growth integrals 10

S1.D.6. Stochastic serial dilution and community lifetime 10

S2. Toxicity as a modulator of temporal growth structure 11

S2.A. The model 11

S2.A.1. Numerical solution of within-cycle dynamics 12

S2.B. Calculating co-growth 12

S2.C. Community lifetime under demographic noise 13

S2.C.1. Deterministic log-ratio map 13

S2.C.2. Stochastic dynamics and definition of lifetime 14

S2.D. Building an ensemble of community "strategies" in the absence of toxicity 14

S2.E. Selecting parameter sets for the toxicity sweep 15

S2.F. Examining the effect of toxicity on co-growth and robustness 15

S3. Extending co-growth as a predictor of robustness to $N > 2$ species 16

S3.A. Extending the invasibility criterion 18

S3.B. Maximum likelihood escape and calculating community lifetime 19

S3.B.1. Defining the minimum action path 19

S3.B.2. First-passage-time representation of the community lifetime 20

S3.B.3. Building an intuition of action as a quasipotential 21

S3.B.4. Linking the escape energy back to invasibility. 22

S4. A phenomenological model of many species 23
S4.A. The model 23
S4.B. Calculating co-growth 24
S4.B.1. Numerical calculation of co-growth (minimum invasibility) 24
S4.C. Community lifetime 25
S4.D. Ensemble construction 26
S4.D.1. Base ensemble construction 26
S4.D.2. Replica-exchange Monte Carlo 27
S4.E. Arrested development upon removal of a single species 28

References 29

### S1. SPATIAL PATTERNING AS A MECHANISM TO STRUCTURE GROWTH IN TIME

Spatially structured communities can maintain coexistence through a variety of mechanisms in which nutrient diffusion, growth, and motility interact to generate self-organized spatial niches [1–5]. In particular, Gude *et al.* [4] showed that coexistence can arise from a trade-off between growth rate and chemotactic ability in colonies expanding on soft agar, inspiring the system we model here. In these experiments, two bacterial strains with different motility and growth rates are inoculated at the center of a circular plate filled with nutrient-rich TB soft agar. The medium contains a primary carbon source that supports growth (e.g., glycerol in TB), as well as amino acids and related compounds present in the rich medium that can serve as chemoattractants. As the strains grow and migrate, they expand radially and typically form a spatially structured pattern, with distinct rings associated with each strain. After hours of incubation, the species abundances were measured. Gude *et al.* then considered a conceptual replating step, in which the population composition after expansion is mapped back to the initial inoculum composition on a new plate. Although this serial-dilution cycle was not performed experimentally, it was introduced as a thought experiment to analyze how species abundances change from one expansion cycle to the next. In this work we explicitly implement serial-dilution *in silico* to study community dynamics over many cycles.

#### S1.A. The model

To describe the spatial growth and expansion dynamics of two species, here, we adopt a mechanistic chemotaxis model from Cremer *et al.* [6]. Two species with densities $\rho_\alpha(r,t)$ ($\alpha = 1,2$) expand on a circular domain of radius $L$ while interacting through a shared nutrient field $n(r,t)$ and chemoattractant field $a(r,t)$.

*a. Species dynamics.* The time evolution of each species' density $\rho_\alpha$ obeys the following partial differential equation:

$$\partial_t \rho_\alpha = D_\rho \nabla^2 \rho_\alpha - \nabla\cdot\big(\mathbf{v}_\alpha \rho_\alpha\big) + \lambda_\alpha(n)\,\rho_\alpha, \qquad \alpha = 1,2, \tag{S1}$$

where the first term represents the passive diffusion with diffusivity $D_\rho$, the second term represents the drift with $\mathbf{v}_\alpha$ via chemotaxis up the chemoattractant gradient, and the last term represents the growth with nutrient-dependent rate $\lambda_\alpha(n)$.

Chemotaxis follows a Weber-law sensing function:

$$\mathbf{v}_\alpha = \chi_{0,\alpha}\,\nabla\left[\log\frac{1+a/a_-}{1+a/a_+}\right], \tag{S2}$$

where $a_-$ and $a_+$ set the receptor sensitivity range and $\chi_{0,\alpha}$ sets the chemotactic strength [6], which is specific to species $\alpha$.

The growth of species follows Monod growth kinetics:

$$\lambda_\alpha(n) = \lambda_{\max,\alpha}\,\frac{n}{n+K_n}. \tag{S3}$$

where $K_n$ is the nutrient half-saturation constant, and $\lambda_{\max,\alpha}$ is a maximum growth rate that is specific to species $\alpha$.

*b. Nutrient dynamics.* Nutrient diffuses with diffusivity $D_n$ and is consumed by growth with yield $Y$:

$$\partial_t n = D_n\,\nabla^2 n - \frac{1}{Y}\sum_{\alpha=1,2}\lambda_\alpha(n)\,\rho_\alpha. \tag{S4}$$

*c. Chemoattractant dynamics.* Chemoattractant diffuses with diffusivity $D_a$ and is taken up by both species at a rate that increases with instantaneous growth and saturates with $a$:

$$\partial_t a = D_a\,\nabla^2 a - \sum_{\alpha=1,2}\mu(\lambda_\alpha(n),a)\;\rho_\alpha, \tag{S5}$$

where

$$\mu(\lambda,a) = \big(c_1\,\lambda + c_2\big)\,\frac{a}{a+K_a}. \tag{S6}$$

Here, $c_1$ couples attractant uptake to growth, $c_2$ captures basal uptake, and $K_a$ is the attractant half-saturation constant.

*d. Rotational symmetry and boundaries.* Note that, assuming rotational symmetry, all fields depend only on the radius $r$ and time $t$. The Laplacian therefore reduces to

$$\nabla^2 f(r) = \frac{1}{r}\,\partial_r(r\,\partial_r f)\,. \tag{S7}$$

Under this symmetry, the model reduces to a one-dimensional radial system. We impose zero-flux (Neumann) boundary conditions at $r=0$ and $r=L$ for all fields.

*e. Traits and parameters.* The focal species-specific traits are the maximal growth rates $\lambda_{\max,\alpha}$ and chemotactic strengths $\chi_{0,\alpha}$; all other parameters are shared and fixed (Table S1). Species 1 traits are fixed to $\lambda_{\max,1} = 0.69$ hr$^{-1}$ and $\chi_{0,1} = 1.13$ mm$^2$/hr. In the ensemble analyzed in the main text, species 2 traits $(\lambda_{\max,2}, \chi_{0,2})$ are sampled on a uniform grid spanning both variables (100 points each, $\lambda_{\max,2} \in [0.5, 1.5]$ 1/hr, $\chi_{0,2} \in (0, 2.5]$ mm$^2$/hr, as reported in the main text), while all non-trait parameters are fixed as in Table S1.

*f. Initial conditions* Initial nutrient and chemoattractant are uniform,

$$n(r,0) = n_0 = 40 \text{ mM}, \qquad a(r,0) = a_0 = 0.1 \text{ mM},$$

and the initial total biomass is a uniform plateau within $r_{\text{plateau}} = 1$ mm,

$$\rho_{\text{tot}}(r,0) = \begin{cases} \rho_0 & r \le r_{\text{plateau}} \\ 0 & r > r_{\text{plateau}} \end{cases}, \tag{S8}$$

with $\rho_0 = 0.029$ OD. Given an initial abundance ratio $X_0 = \tilde{\rho}_1/\tilde{\rho}_2$, we set

$$\rho_1(r,0) = \frac{X_0}{1+X_0}\rho_{\text{tot}}(r,0), \qquad \rho_2(r,0) = \frac{1}{1+X_0}\rho_{\text{tot}}(r,0).$$

Each cycle is integrated from $t = 0$ to $t = T_{\max}$ and the end-of-cycle ratio $X_T = \tilde{\rho}_1(T_{\max})/\tilde{\rho}_2(T_{\max})$ is recorded, where $\tilde{\rho}_\alpha(t)$ is the spatially integrated biomass of species $\alpha$ is

$$\tilde{\rho}_\alpha(t) = 2\pi \int_0^L r\, \rho_\alpha(r,t)\, dr.$$

#### S1.B. Calculating co-growth with spatial heterogeneity

Integrating Eqn. (S1) over the plate and applying the divergence theorem shows that diffusion and chemotactic advection contribute only via boundary fluxes. With zero-flux (Neumann) boundary conditions at $r = 0$ and $r = L$, these boundary terms vanish, so total biomass changes only through growth:

$$\frac{d\tilde{\rho}_\alpha}{dt} = \tilde{\lambda}_\alpha(t)\, \tilde{\rho}_\alpha(t),$$

with global per-capita growth rate

$$\tilde{\lambda}_\alpha(t) = \frac{2\pi}{\tilde{\rho}_\alpha(t)} \int_0^L r\, \lambda_\alpha(n(r,t))\, \rho_\alpha(r,t)\, dr.$$

TABLE S1. Model parameters for the spatial patterning model (Section S1). Parameter values match those used in Cremer *et al.* [6], except for the plate radius $L$ (set to match [4]) and the species-2 traits ($\lambda_{\mathrm{max},2}, \chi_{0,2}$), which are varied as described in the text. Unless otherwise noted, parameters are shared by both species; species-specific traits are $\lambda_{\mathrm{max},\alpha}$ and $\chi_{0,\alpha}$.

| **Parameter** | **Description** | **Value** | **Notes** |
|---|---|---|---|
| $L$ | Plate radius | 17.5 mm | Fixed |
| $D_\rho$ | Cell diffusivity | 0.18 $\mathrm{mm}^2/\mathrm{hr}$ | Same for both species |
| $D_n$ | Nutrient diffusivity | 2.88 $\mathrm{mm}^2/\mathrm{hr}$ | Fixed |
| $D_a$ | Chemoattractant diffusivity | 2.88 $\mathrm{mm}^2/\mathrm{hr}$ | Fixed |
| $Y$ | Growth yield | 0.064 OD/mM | Fixed |
| $\chi_{0,1}$ | Chemotactic strength, species 1 | 1.13 $\mathrm{mm}^2/\mathrm{hr}$ | Fixed |
| $\chi_{0,2}$ | Chemotactic strength, species 2 | | Varied in ensemble |
| $a_-$ | Weber-law lower threshold | 0.001 mM | Fixed |
| $a_+$ | Weber-law upper threshold | 0.03 mM | Fixed |
| $\lambda_{\mathrm{max},1}$ | Max growth rate, species 1 | 0.69 $\mathrm{hr}^{-1}$ | Fixed |
| $\lambda_{\mathrm{max},2}$ | Max growth rate, species 2 | | Varied in ensemble |
| $K_n$ | Nutrient half-saturation (Monod) | 0.05 mM | Fixed |
| $c_1$ | Attractant uptake (growth-coupled) | 0.77 mM/(OD·hr) | Fixed |
| $c_2$ | Attractant uptake (basal) | 1.25 mM/OD | Fixed |
| $K_a$ | Attractant half-saturation (Monod) | 0.001 mM | Fixed |
| $n_0$ | Initial nutrient concentration | 40 mM | Uniform across plate |
| $a_0$ | Initial chemoattractant concentration | 0.1 mM | Uniform across plate |
| $\rho_0$ | Initial biomass density | 0.029 OD | In inoculation zone |
| $r_{\mathrm{plateau}}$ | Inoculation radius | 1.0 mm | Fixed |
| $T_{\mathrm{max}}$ | Duration of one growth cycle | 24 hr | Fixed |

See Fig. 2D for examples of the time dependence of $\tilde{\lambda}_\alpha(t)$.

Using $\tilde{\lambda}_\alpha(t)$ in the co-growth framework (Eqn. 5 in the main text), we compute

$$I_{12} = \int_0^{T_{\mathrm{max}}} \frac{\tilde{\lambda}_1}{\tilde{\lambda}_2} \frac{d \ln \tilde{\rho}_{\mathrm{tot}}}{dt}\, dt, \qquad I_{21} = \int_0^{T_{\mathrm{max}}} \frac{\tilde{\lambda}_2}{\tilde{\lambda}_1} \frac{d \ln \tilde{\rho}_{\mathrm{tot}}}{dt}\, dt, \tag{S9}$$

where $\tilde{\rho}_{\mathrm{tot}} = \tilde{\rho}_1 + \tilde{\rho}_2$, and define the co-growth metric

$$I = \min(I_{12},\ I_{21})\,. \tag{S10}$$

Note that the values of $I_{12}$ and $I_{21}$ depend on the initial abundance ratio $X_0$; we set $X_0 = X_{\mathrm{extinct}}$ or $X_0 = 1/X_{\mathrm{extinct}}$ to calculate $I_{12}$ or $I_{21}$, respectively.

#### S1.C. Community lifetime under demographic noise

We next consider the dispersal cycle, in which the species abundance ratio $X = \tilde{\rho}_1/\tilde{\rho}_2$ is transferred between successive growth cycles. Note that the dilution ratio $\delta$ in Eqn. 4 of the main text is, in this spatial system, given by $\delta = \tilde{\rho}^{\mathrm{tot}}(0)/\tilde{\rho}^{\mathrm{tot}}(T_{\max})$, where the initial total biomass is $\tilde{\rho}^{\mathrm{tot}}(0) = \pi(r_{\mathrm{plateau}})^2\rho_0$, and one at the end-of-cycle is $\tilde{\rho}^{\mathrm{tot}}(T_{\max}) = \pi(r_{\mathrm{plateau}})^2\rho_0 + \pi L^2 n_0$, assuming that all nutrients are consumed by the end of the cycle.

We consider parameter sets $\{\chi_{0,\alpha}, \lambda_{\max,\alpha}\}_{\alpha=1,2}$ for which a deterministic fixed point of the species abundance ratio $X^*$ exists.

The stochastic dispersal cycles are defined as a map

$$\log X^{(j+1)} = \log X^{(j)} + \Delta \log X^{(j)} + \sqrt{\sigma}\,\xi^{(j)}, \tag{S11}$$

where $\xi^{(j)} \sim \mathcal{N}(0,1)$ and $\sigma$ sets the noise strength, and $\Delta \log X^{(j)}$ is the change in log species abundance ratio as a function of the ratio at the start of the cycle, defined by the growth-expansion dynamics described previously. The stochastic term represents demographic noise during dispersal.

We define the lifetime of the community as the first passage time until the ratio $X$ becomes below or exceeds the extinction threshold, $X_{\mathrm{extinct}}$ or $1/X_{\mathrm{extinct}}$, starting from $X^{(0)} = X^*$.

#### S1.D. Numerical implementation and ensemble construction

We solved Eqs. (S1)–(S6) using an explicit finite-difference scheme in radial coordinates (C++; rotational symmetry).

*S1.D.1. Spatial discretization and boundary conditions*

The domain is a disk of radius $L = 17.5$ mm, discretized into $N_r = 100$ concentric annuli with uniform spacing $\Delta r = L/N_r$. Grid points are cell-centered,

$$r_i = \left(i + \tfrac{1}{2}\right)\Delta r, \qquad i = 0, \ldots, N_r - 1.$$

We discretize divergence-form radial operators using face fluxes. For terms of the form $\frac{1}{r}\partial_r(rJ)$,

$$\frac{1}{r}\frac{\partial}{\partial r}(rJ)\bigg|_{r=r_i} \approx \frac{r_{i+1/2}\, J_{i+1/2} - r_{i-1/2}\, J_{i-1/2}}{r_i\, \Delta r},$$

with $r_{i\pm1/2} = r_i \pm \Delta r/2$ and face values formed from neighboring cell-center values (arithmetic averages for densities and concentrations).

For chemotaxis, the face velocity uses the Weber-law prefactor evaluated at the face-averaged attractant:

$$\left[\frac{1}{a_- + a} - \frac{1}{a_+ + a}\right]\frac{\partial a}{\partial r}\bigg|_{i+1/2} \approx \left[\frac{1}{a_- + \bar{a}_{i+1/2}} - \frac{1}{a_+ + \bar{a}_{i+1/2}}\right]\frac{a_{i+1} - a_i}{\Delta r}, \qquad \bar{a}_{i+1/2} = \frac{a_i + a_{i+1}}{2},$$

and we use $\rho_{i+1/2} = (\rho_i + \rho_{i+1})/2$ for the advected density at the face.

Zero-flux boundary conditions at $r = 0$ and $r = L$ are imposed via reflecting ghost values (equivalently enforcing a discrete Neumann condition):

$$y_0 = y_1, \qquad y_{N_r-1} = y_{N_r-2},$$

for each field $y \in \{\rho_1, \rho_2, n, a\}$.

Spatial integrals use the midpoint rule,

$$2\pi \int_0^L r\, f(r)\, dr \approx \sum_{i=0}^{N_r-1} 2\pi\, r_i\, f(r_i)\, \Delta r.$$

*S1.D.2. Time integration within one growth cycle*

Time evolution uses forward Euler,

$$y(r_i, t + \Delta t) = y(r_i, t) + \Delta t\, \partial_t y(r_i, t),$$

with time step $\Delta t = 1$ s ($= 1/3600$ hr) and cycle duration $T_{\max} = 24$ hr.

*S1.D.3. Identification of coexistence parameter sets*

We compute $\Delta \log X = \log X_T - \log X_0$ for two extreme initial ratios, $\log X_0 = \pm \log X_{\text{extinct}}$ with $\log X_{\text{extinct}} = 10$. Parameter sets satisfying

$$\Delta \log X(\log X_0 = -10) \ \cdot \ \Delta \log X(\log X_0 = +10) < 0$$

are retained, indicating a sign change and therefore a stable coexistence fixed point.

*S1.D.4. Constructing the $\Delta \log X$ map*

For each coexistence-eligible parameter set, we simulate one growth cycle over a grid of initial ratios spanning $\log X \in [-10, 10]$ with step size 0.5, recording $\Delta \log X$ at each grid point. This map is then used to iterate the serial-dilution dynamics efficiently over many cycles.

*S1.D.5. Co-growth integrals*

The co-growth integrals in Eqn. (S9) are evaluated as discrete sums over time steps within a single-cycle simulation:

$$I_{12} \approx \sum_{\ell} \frac{\tilde{\lambda}_1(t_\ell)}{\tilde{\lambda}_2(t_\ell)} \frac{\rho_{\text{tot}}(t_\ell) - \rho_{\text{tot}}(t_{\ell-1})}{\rho_{\text{tot}}(t_\ell)}, \qquad I_{21} \approx \sum_{\ell} \frac{\tilde{\lambda}_2(t_\ell)}{\tilde{\lambda}_1(t_\ell)} \frac{\rho_{\text{tot}}(t_\ell) - \rho_{\text{tot}}(t_{\ell-1})}{\rho_{\text{tot}}(t_\ell)}.$$

To probe each invasion limit, we compute $I_{12}$ from a cycle initialized at $X_0 = X_{\text{extinct}}^{-1}$ (species 1 rare) and $I_{21}$ from a cycle initialized at $X_0 = X_{\text{extinct}}$ (species 2 rare).

*S1.D.6. Stochastic serial dilution and community lifetime*

Deterministic fixed points $X^*$ are estimated as the $\log X$ value where $|\Delta \log X|$ is minimized in the precomputed map. Starting from $\log X^{(0)} = \log X^*$, we iterate a stochastic map

$$\log X^{(j+1)} = \log X^{(j)} + \Delta \log X^{(j)} + \sqrt{\sigma}\, \xi^{(j)},$$

where $\xi^{(j)} \sim \mathcal{N}(0, 1)$ and $\sigma$ sets the noise strength. Values of $\Delta \log X$ at off-grid $\log X$ are obtained by linear interpolation.

A community is considered extinct when $|\log X^{(j)}| > \log X_{\mathrm{extinct}} = 10$. If extinction does not occur within a specified maximum number of cycles, the lifetime is recorded as $\infty$. For each parameter set we generate 100 independent stochastic trajectories and compute the mean lifetime over finite trajectories ($\sigma = 4.5$); together with the corresponding co-growth metrics, these define the ensemble analyzed in Fig. 2.

### S2. TOXICITY AS A MODULATOR OF TEMPORAL GROWTH STRUCTURE

#### S2.A. The model

The dynamics of the two species (Nar$^+$, hereafter species 1, $\rho_1(t)$; Nap$^+$, hereafter species 2, $\rho_2(t)$) and two metabolites ($NO_3^-$: $n_A$, $NO_2^-$: $n_I$) within each growth cycle were calculated following a numerical model inspired by [7, 8]. It is assumed that the system is well-mixed.

A community strategy is defined by the two species' resource-specific maximum growth rates on nitrate and nitrite, $\{\mu_{\alpha,A},\ \mu_{\alpha,I}\}_{\alpha=1,2}$, while yield ($\gamma = 1$), the half-saturation constants ($K_A = K_I = 0.01$, $K_g = 0.5\, n_{A,0} = 5$), and the Hill coefficient for the gain function ($h = 10$) are held constant. Toxicity is introduced as a graded parameter between 0 (no toxicity) and 1 (maximal toxicity).

$$\frac{d\rho_1}{dt} = g_1(n_I)\left(\gamma\,\mu_{1,A}\frac{n_A}{K_A+n_A} + \gamma\,\mu_{1,I}\frac{n_I}{K_I+n_I}\right)\rho_1 \tag{S12}$$

$$\frac{d\rho_2}{dt} = \left(\gamma\,\mu_{2,A}\frac{n_A}{K_A+n_A} + \gamma\,\mu_{2,I}\frac{n_I}{K_I+n_I}\right)\rho_2 \tag{S13}$$

$$\frac{dn_A}{dt} = -\big(g_1(n_I)\,\mu_{1,A}\,\rho_1 + \mu_{2,A}\,\rho_2\big)\frac{n_A}{K_A+n_A} \tag{S14}$$

$$\frac{dn_I}{dt} = -\frac{dn_A}{dt} - \big(g_1(n_I)\,\mu_{1,I}\,\rho_1 + \mu_{2,I}\,\rho_2\big)\frac{n_I}{K_I+n_I} \tag{S15}$$

$$g_1(n_I) = 1 - \frac{\text{toxicity}}{1 + \big(K_g/(n_I+\varepsilon)\big)^h} \tag{S16}$$

Only species 1 (Nar$^+$) is affected by toxicity; species 2 (Nap$^+$) is not. The small regularization $\varepsilon = \varepsilon_{\mathrm{mach}} \approx 2.2 \times 10^{-16}$ is added to the denominator of the gain function to avoid a singularity at $n_I = 0$; it ensures that $g_1(0) = 1$ regardless of toxicity, consistent with the biological expectation that the inhibitory effect of nitrite is absent at the start of the cycle when no nitrite has yet accumulated.

The initial nutrient levels are fixed ($n_A(0) = n_{A,0} = 10$ and $n_I(0) = 0$), so nitrite is produced only through the metabolism of nitrate. The initial total biomass is fixed at

$\rho_{\text{tot},0} = 1$ and the initial species ratio $X = \rho_1(0)/\rho_2(0)$ is varied. The total biomass at the end of a cycle is $\rho_{\text{tot}}^{\max} = 2n_{A,0} + \rho_{\text{tot},0} = 21$: nitrate is fully converted to nitrite ($\gamma = 1$, stoichiometry 1:1 for N), and all nitrite is in turn consumed to produce biomass, so total final biomass $= 2n_{A,0} + \rho_{\text{tot},0}$. The dilution fraction is therefore $\delta = \rho_{\text{tot},0}/\rho_{\text{tot}}^{\max} = 1/21$; multiplying by $\delta$ restores the initial total biomass to $\rho_{\text{tot},0} = 1$ at the start of each cycle.

##### *S2.A.1. Numerical solution of within-cycle dynamics*

For a given community, specified by its growth rates $\{\mu_{\alpha,A},\ \mu_{\alpha,I}\}_{\alpha=1,2}$ and a particular toxicity level, we numerically integrated the ODEs in SI Sec. S2.A using a standard adaptive-step explicit Runge–Kutta method (Dormand–Prince, `RK45` in `solve_ivp`, SciPy). The state vector was

$$y(t) = \big(n_A(t),\, n_I(t),\, \rho_1(t),\, \rho_2(t)\big),$$

and the right-hand side implemented Eqs. (S12)–(S16). All cycles used the same initial conditions:

$$n_A(0) = n_{A,0} = 10, \qquad n_I(0) = 0, \qquad \rho_1(0) + \rho_2(0) = \rho_{\text{tot},0} = 1,$$

with $X = \rho_1(0)/\rho_2(0)$ setting the initial community composition. Non-negativity of all state variables was enforced by truncating any numerically negative values to zero at each evaluation of the right-hand side.

Integration was halted when both resources were depleted:

$$\max\{n_A(t),\, n_I(t)\} \le 10^{-12}.$$

#### S2.B. Calculating co-growth

For a given community, we quantified co-growth using the integrals $I_{12}$ and $I_{21}$ defined in the main text (Eqn. 5):

$$I_{12} = \int \frac{\lambda_1(\rho_{\text{tot}})}{\lambda_2(\rho_{\text{tot}})} \frac{d\rho_{\text{tot}}}{\rho_{\text{tot}}}, \qquad I_{21} = \int \frac{\lambda_2(\rho_{\text{tot}})}{\lambda_1(\rho_{\text{tot}})} \frac{d\rho_{\text{tot}}}{\rho_{\text{tot}}},$$

where $\lambda_\alpha$ is the instantaneous per-capita growth rate of species $\alpha$ along a growth cycle and $\rho_{\text{tot}} = \rho_1 + \rho_2$ is the total biomass. Numerically, we obtained $\lambda_\alpha$ from the ODE solution via

$$\lambda_1(t) = \frac{1}{\rho_1(t)} \frac{d\rho_1}{dt}, \qquad \lambda_2(t) = \frac{1}{\rho_2(t)} \frac{d\rho_2}{dt},$$

evaluated along the trajectory from $(n_A(0), n_I(0)) = (n_{A,0}, 0)$ until resource depletion, and approximated the integrals over $\rho_{\text{tot}}$ by trapezoidal quadrature.

Each invasion scenario was initialised near monoculture: $I_{12}$ used $(\rho_1(0), \rho_2(0)) = (\varepsilon_{\text{mach}},\, 1 - \varepsilon_{\text{mach}})$ and $I_{21}$ used $(\rho_1(0), \rho_2(0)) = (1 - \varepsilon_{\text{mach}},\, \varepsilon_{\text{mach}})$, where $\varepsilon_{\text{mach}} \approx 2.2 \times 10^{-16}$ is machine epsilon.

Communities were classified as coexisting when co-growth exceeded (following [9]):

$$\min(I_{12}, I_{21}) > -\log \delta,$$

where $-\log \delta = \log(21) \approx 3.04$.

#### S2.C. Community lifetime under demographic noise

To quantify robustness, we defined the community lifetime as the expected number of growth-dilution cycles until one species is lost ($|\log X| > \log X_{\text{extinct}}$, where $\log X_{\text{extinct}} = 10$). This calculation uses a one-dimensional map for the species log ratio, $\Delta \log X$, where $X = \rho_1/\rho_2$.

##### *S2.C.1. Deterministic log-ratio map*

For each parameter set we reduced the full consumer-resource dynamics to a deterministic per-cycle map for $\log X$ at fixed total initial biomass $\rho_{\text{tot}}(0) = 1$. For a grid of 101 evenly spaced initial log ratios,

$$\log X \in [-(\log X_{\text{extinct}} + 1),\, +(\log X_{\text{extinct}} + 1)],$$

we set

$$\rho_1(0) = \frac{X}{1+X}, \qquad \rho_2(0) = \frac{1}{1+X},$$

with $n_A(0) = n_{A,0}$, $n_I(0) = 0$, and numerically integrated through one full growth cycle using the solver described in SI Sec. S2.A.1. The deterministic change across the cycle is

$$\Delta \log X \equiv \log X' - \log X,$$

yielding a lookup table $\{\log X_i,\, \Delta \log X_i\}_{i=1}^{101}$.

A coexistence steady state is identified as the grid point $\log X^*$ where $|\Delta \log X|$ is minimised, subject to the requirement that $\Delta \log X$ passes through zero from positive to negative (a stable interior fixed point not at the grid boundaries). Parameter sets that failed this condition were excluded from the lifetime analysis.

#### *S2.C.2. Stochastic dynamics and definition of lifetime*

Starting from the deterministic fixed point $\log X^{(0)} = \log X^*$, we iterated the stochastic map

$$\log X^{(j+1)} = \log X^{(j)} + \Delta \log X^{(j)} + \sqrt{\sigma}\, \xi^{(j)},$$

where $\Delta \log X^{(j)}$ is obtained by linear interpolation of the precomputed lookup table, $\xi^{(j)} \sim \mathcal{N}(0, 1)$ is an independent Gaussian increment, and $\sigma$ is the demographic noise amplitude (variance per cycle). A community is considered extinct when $|\log X^{(j)}| > \log X_{\mathrm{extinct}} = 10$. We ran 1000 replicates per parameter set (with $\sigma = 2.75$) and recorded the mean first-passage time across replicates that went extinct within $10^6$ cycles; replicates that did not go extinct within this limit were recorded as $\tau = \infty$, and points for which any replicate had $\tau = \infty$ were excluded.

### S2.D. Building an ensemble of community "strategies" in the absence of toxicity

To systematically map the co-growth landscape for denitrifying communities, we fixed species 1 at an anchor point $(\mu_{1,A}, \mu_{1,I})$ and swept the species 2 growth rates $(\mu_{2,A}, \mu_{2,I})$ over a $100 \times 100$ grid:

$$\mu_{2,A},\ \mu_{2,I} \in [0,\ 1.5], \quad \text{100 evenly spaced points per axis},$$

giving 10,000 parameter sets in total. For each grid point we computed $I_{12}$, $I_{21}$, and the coexistence flag ($\min(I_{12}, I_{21}) > -\log \delta$). An advantage label was assigned based on which invasibility metric exceeded the other: $I_{12} > I_{21}$ (species 1 advantage) or $I_{12} < I_{21}$ (species 2 advantage).

Two anchor points were examined:

- **Anchor 01**: $\mu_{1,A} = 0.710$, $\mu_{1,I} = 0.093$ — chosen because, at this anchor, toxicity had both a *beneficial* and *detrimenal* effect on robustness.

- **Anchor 02**: $\mu_{1,A} = 0.377$, $\mu_{1,I} = 0.692$ — chosen as a contrast to anchor 01; overall coexistence region of parameter space in the absence of toxicity is similar, but now the toxicity effect on robustness is *neutral* or *detrimental.*

The results of each base sweep can be seen in Figs. S1a and S2a.

#### S2.E. Selecting parameter sets for the toxicity sweep

From the base sweep we retained only coexisting parameter sets and stratified them by advantage. Within each advantage class, we further divided by cone, defined relative to the anchor nitrite growth rate $\mu_{1,I}$:

- **Top cone**: $\mu_{2,I} > \mu_{1,I}$ (species 2 has a higher nitrite growth rate than species 1),
- **Bottom cone**: $\mu_{2,I} \leq \mu_{1,I}$ (species 2 has a lower nitrite growth rate than species 1).

Within each advantage-cone combination, parameter sets were sampled randomly, with the total budget split as evenly as possible between the two cones. This stratification captures qualitatively distinct competitive contexts: in the top cone, species 2 can exploit the nitrite that accumulates during the cycle more aggressively than species 1 whereas in the bottom cone the two species are more comparable in their nitrite kinetics.

#### S2.F. Examining the effect of toxicity on co-growth and robustness

For each selected parameter set we swept toxicity $\in [0, 1]$ at 101 uniformly spaced values and recomputed $I_{12}$, $I_{21}$, and $I = \min(I_{12}, I_{21})$ at each level. The resulting traces (Fig. S1b and Fig. S2b) reveal that the direction of toxicity's effect on robustness depends on which species holds the competitive advantage.

When Species 2 holds the advantage ($I_{21} > I_{12}$), toxicity is either neutral, leaving $I = \min(I_{12}, I_{21})$, co-growth, and community lifetime unchanged, or detrimental, decreasing $I = \min(I_{12}, I_{21})$, increasing co-growth, and reducing robustness. When Species 1 holds the advantage ($I_{12} > I_{21}$), toxicity is either again neutral or, in contrast, beneficial: $I = \min(I_{12}, I_{21})$ increases, co-growth decreases, and robustness improves.

These outcomes are layered on top of an already complex parameter landscape shaped by growth rates on the two resources (nitrate and nitrite), making it exceedingly difficult to predict *a priori* whether toxicity will exert a beneficial, detrimental, or neutral influence in any given region of parameter space; additional regimes with yet other qualitative effects may as well exist.

The key point for this manuscript, however, is that irrespective of this complexity and the direction of toxicity's effect, co-growth continues to serve as a reliable predictor of robustness. To demonstrate this, in the main text (Fig. 3c) we highlight two traces from those shown here (starred points in Fig. S1a; darker traces in Fig. S1b), representing two entirely different toxicity-co-growth trends, and calculate the community lifetime (see SI Sec. S2.C) to show that in both cases robustness tracks co-growth in a manner consistent with our log-linear expectation (see main text Sec. IV).

### S3. EXTENDING CO-GROWTH AS A PREDICTOR OF ROBUSTNESS TO $N > 2$ SPECIES

When extending this theory to $N > 2$ species, it helps to first build an intuition for the argument.

In two species, the dynamics can be described on a 1D line of log-ratio of abundances and we were able to construct an quasi potential on that line. From this quasi potential we could calculate the expected community lifetime with Arrhenius's Law, which is a function of the escape energy. This escape energy built the link between expected community lifetime and co-growth between the two species by the observation that the potential energy at the extinction boundary of one species (e.g species 1) is approximately proportional to the invasibility factor ($I_{12}$) of that species.

With $N > 2$ species, the dynamics now lie on the $N - 1$ dimensional simplex describing relative abundance of the $N$ species. While it is no longer possible to generally construct an quasi potential over the whole space of dynamics, because we cannot guarantee that the vector field is curl-free, we can still (i) examine the existence of a stable point with full coexistence using the vector field along the boundary of the simplex and (ii) examine the time to escape by considering the escape trajectory of greatest likelihood.

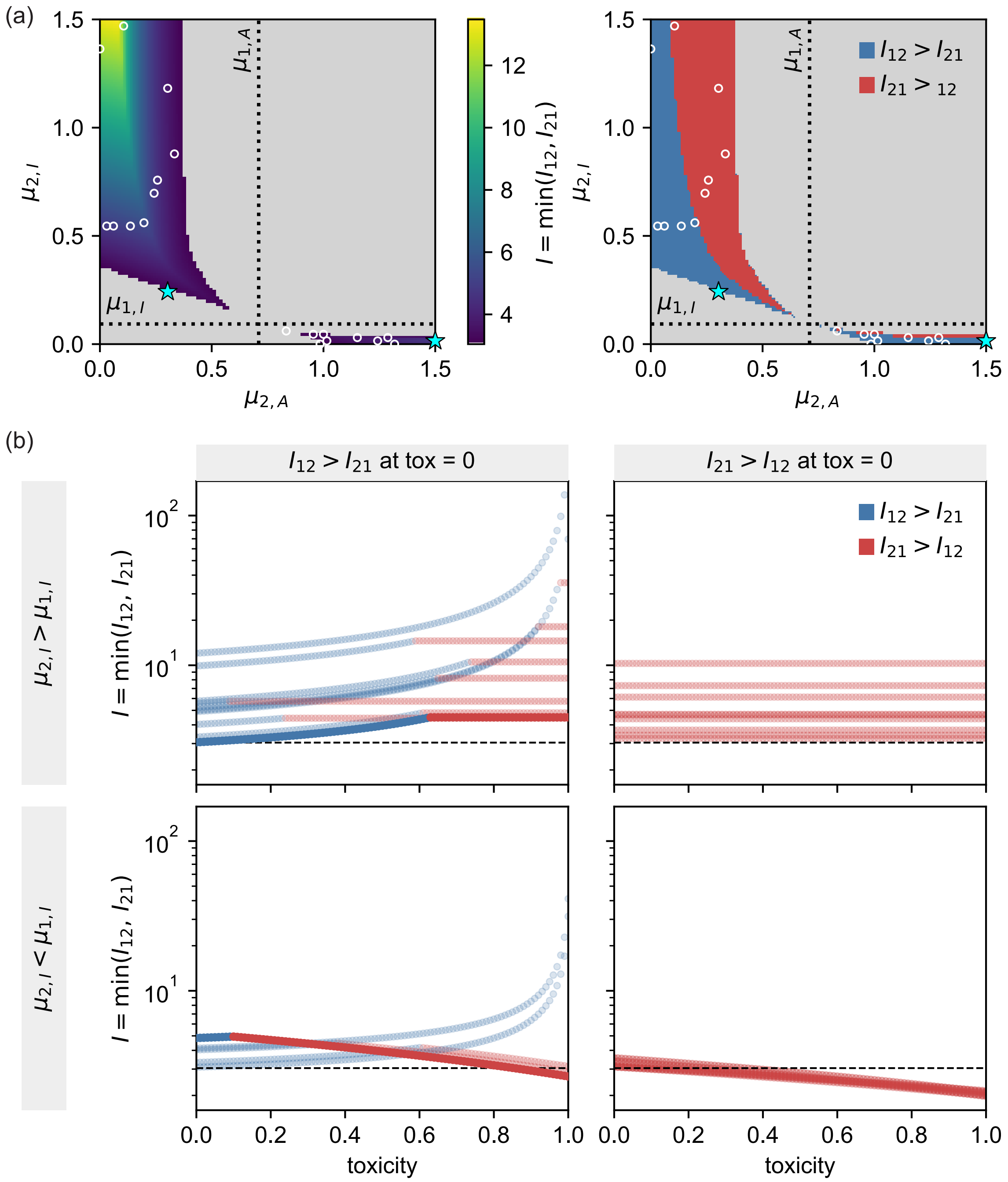


FIG. S1. **Parameter space exploration and toxicity sweep for Anchor 01 ($\mu_{1,A} = 0.710$, $\mu_{1,I} = 0.093$).** **(a)** Heatmaps over species 2 growth rates $(\mu_{2,A}, \mu_{2,I})$ at zero toxicity: $I = \min(I_{12}, I_{21})$ (left) and advantage map (right; blue: species 1, $I_{12} > I_{21}$; red: species 2, $I_{21} > I_{12}$). Grey denotes no-coexistence regions. Dashed lines mark the anchor; scatter points indicate parameter sets selected for the toxicity sweep (star: main-text cases; circles: SI only). **(b)** $I = \min(I_{12}, I_{21})$ as a function of toxicity with points colored by instantaneous advantage. Darker traces correspond to those plotted in the main text (Fig. 3c). Dashed line: coexistence threshold $(-\log \delta)$.

#### S3.A. Extending the invasibility criterion

First we extend the invasibility criterion to $N > 2$ species by considering the vector field around the perimeter of the simplex, following a simlar derivation to the one put forth in [9]. Suppose we have a system initialized with abundances $\boldsymbol{\rho}_0^{\setminus\alpha}$, where $\rho_{\alpha,0}^{\setminus\alpha} = \epsilon$ and $\rho_{\beta,0}^{\setminus\alpha} \geq 0$ for all $\beta \neq \alpha$, with $\epsilon \ll \rho_0 = \sum_{\beta\neq\alpha} \rho_{\beta,0}^{\setminus\alpha}$ the initial biomass of the subcommunity. In other words, we examine the invasion of a rare species $\alpha$ into an arbitrary subcommunity of the remaining species; we do not require that all $N-1$ other species be present. We can then write out the log-fold increase of species $\alpha$

$$\log\left(\frac{\rho_\alpha^{(j+1)}(0)}{\rho_\alpha^{(j)}(0)}\right) = \int_0^T \lambda_\alpha(\rho_{\text{tot}}(t))dt + \log(\delta). \tag{S17}$$

Provided $\rho_{\text{tot}}(t)$ is monotonic in time, we can do a variable substitution

$$\begin{aligned} \log\left(\frac{\rho_\alpha^{(j+1)}(0)}{\rho_\alpha^{(j)}(0)}\right) &= \int_{\rho^0+\epsilon}^{\rho_{\text{tot}}^{\max}} \frac{\lambda_\alpha(\rho_{\text{tot}})}{d\rho_{\text{tot}}/dt} d\rho_{\text{tot}} + \log(\delta) \\ &= \int_{\rho^0+\epsilon}^{\rho_{\text{tot}}^{\max}} \frac{\lambda_\alpha(\rho_{\text{tot}})}{\sum_\beta \lambda_\beta(\rho_{\text{tot}})\rho_\beta(\rho_{\text{tot}})} d\rho_{\text{tot}} + \log(\delta) \end{aligned} \tag{S18}$$

Now assuming that $\lambda_\beta(\rho_{\text{tot}}) > 0, \forall\, \beta$ and $\rho_{\text{tot}}$, we can always choose an $\epsilon$ such that $\rho_\alpha(\rho_{\text{tot}}) \ll \rho_\beta(\rho_{\text{tot}})\ \forall\ \beta \neq \alpha$ and $\rho_{\text{tot}}$ and thus $\lambda_\alpha(\rho_{\text{tot}})\rho_\alpha(\rho_{\text{tot}}) \ll \lambda_\beta(\rho_{\text{tot}})\rho_\beta(\rho_{\text{tot}})\ \forall\ \beta \neq \alpha$ and $\rho_{\text{tot}}$, yielding the following

$$\log\left(\frac{\rho_\alpha^{(j+1)}(0)}{\rho_\alpha^{(j)}(0)}\right) \approx \int_{\rho^0}^{\rho_{\text{tot}}^{\max}} \frac{\lambda_\alpha(\rho_{\text{tot}})}{\sum_{\beta\neq\alpha} \lambda_\beta(\rho_{\text{tot}})\rho_\beta(\rho_{\text{tot}})} d\rho_{\text{tot}} + \log(\delta). \tag{S19}$$

The structure of this integral is very similar to the $I_{12}$, but now instead of having a simple ratio of the two growth rates ($\lambda_1/\lambda_2$) we have a ratio of the growth rate of the invading species, $\lambda_\alpha(\rho_{\text{tot}})$, compared against the weighted sum of the other species ($\sum_{\beta\neq\alpha} \lambda_\beta(\rho_{\text{tot}})\rho_\beta(\rho_{\text{tot}})$). This new ratio can be interpreted as measuring the growth advantage that one species has against the community average growth rate; again this value should be small when there is considerable coincidental growth and larger when there is more separation of growth. Note that this integral is a function of the individual abundances via the weighted average growth rate in the denominator, which means that the gradient away from the boundary will be dependent on where you are along the boundary. In the case of $N = 2$ species, invasion is more simply defined because a species can invade only into a monoculture of the other species. In the case of $N > 2$, invasion is now defined on a simplex of $N - 2$ dimensions, so the starting composition of that sub-community becomes relevant even if the total biomass is fixed.

To assess invasibility, we look at the change over many cycles. In subsequent cycles, the log-fold increase of species $\alpha$ will be

$$
\begin{aligned}
\log\left(\frac{\rho_\alpha^{(j+k+1)}(0)}{\rho_\alpha^{(j+k)}(0)}\right) &\approx \int_{\delta\rho_{\rm tot}^{\max}}^{\rho_{\rm tot}^{\max}} \frac{\lambda_\alpha(\rho_{\rm tot})}{\sum_{\beta\neq\alpha}\lambda_\beta(\rho_{\rm tot})\rho_\beta(\rho_{\rm tot})} d\rho_{\rm tot} + \log(\delta) \\
\log\left(\frac{\rho_\alpha^{(j+k+1)}(0)}{\rho_\alpha^{(j)}(0)}\right) &\approx k\int_{\delta\rho_{\rm tot}^{\max}}^{\rho_{\rm tot}^{\max}} \frac{\lambda_\alpha(\rho_{\rm tot})}{\sum_{\beta\neq\alpha}\lambda_\beta(\rho_{\rm tot})\rho_\beta(\rho_{\rm tot})} d\rho_{\rm tot} \\
&\quad + \int_{\rho^0}^{\rho_{\rm tot}^{\max}} \frac{\lambda_\alpha(\rho_{\rm tot})}{\sum_{\beta\neq\alpha}\lambda_\beta(\rho_{\rm tot})\rho_\beta(\rho_{\rm tot})} d\rho_{\rm tot} \\
&\quad + (k+1)\log(\delta)
\end{aligned}
\tag{S20}
$$

which means that in order for species $\alpha$ to invade, you need to satisfy for $k \to \infty$ that $\rho_\alpha^{(j+k+1)}(0) \gg \epsilon$, in other words that

$$
I_\alpha \equiv \int_{\delta\rho_{\rm tot}^{\max}}^{\rho_{\rm tot}^{\max}} \frac{\lambda_\alpha(\rho_{\rm tot})}{\sum_{\beta\neq\alpha}\lambda_\beta(\rho_{\rm tot})\rho_\beta(\rho_{\rm tot})} d\rho_{\rm tot} > -\log(\delta) \tag{S21}
$$

for all community compositions within the $N-1$ species subspace. The strict condition for coexistence is therefore that $I_\alpha(\boldsymbol{\rho}_0^{\backslash\alpha}) > -\log(\delta)$ for all species $\alpha$ and all sub-community compositions $\boldsymbol{\rho}_0^{\backslash\alpha}$, or equivalently $\min_{\alpha,\boldsymbol{\rho}_0^{\backslash\alpha}} I_\alpha(\boldsymbol{\rho}_0^{\backslash\alpha}) > -\log(\delta)$. This is the natural $N$-species generalization of the two-species condition $\min(I_{12}, I_{21}) > -\log(\delta)$: the minimization now runs over both all invasion directions *and* all possible compositions along the boundary of the simplex.

#### S3.B. Maximum likelihood escape and calculating community lifetime

Just like before in the case of $N = 2$ species, the average community lifetime relates to rare extinction events using a first-passage-time formulation, however in the case of $N > 2$ species the different paths that the system can take result in escaping from different "exits".

##### *S3.B.1. Defining the minimum action path*

Recall that we model cycle-to-cycle fluctuations in the initial density of species $\alpha$ by the discrete map

$$
\log\rho_\alpha^{(j+1)}(0) = \log\rho_\alpha^{(j)}(0) + \Delta\log\rho_\alpha^{(j)} + \sqrt{\sigma/2}\,\xi_\alpha^{(j)}, \tag{S22}
$$

where $\xi_\alpha^{(j)} \sim \mathcal{N}(0,1)$ are i.i.d. Gaussian noises, $\sigma$ sets the noise strength, and $\Delta \log \rho_\alpha^{(j)}$ captures the deterministic dynamics during one growth cycle. For clarity, we work with an effective per-species description in which log-density increments are treated as independent Gaussian noise terms with amplitude $\sqrt{\sigma/2}$; this matches the per-species amplitude of the physical model (Eqn. 4) while neglecting the total-density conservation constraint that couples species through $Z^{(j)}$.

Rearranging, the realized noise at cycle $j$ is the residual between the observed change in log-density and the deterministic drift:

$$\xi_\alpha^{(j)} = \frac{\log \rho_\alpha^{(j+1)}(0) - \log \rho_\alpha^{(j)}(0) - \Delta \log \rho_\alpha^{(j)}}{\sqrt{\sigma/2}}. \tag{S23}$$

The probability weight of a trajectory $\{\log \boldsymbol{\rho}^{(0)}, \ldots, \log \boldsymbol{\rho}^{(k)}\}$ is therefore (by the Onsager-Machlup construction for Gaussian increments [10, 11])

$$p(\log \boldsymbol{\rho}^{(0)}, \ldots, \log \boldsymbol{\rho}^{(k)}) \propto \prod_{j=0}^{k-1} \prod_{\alpha} \exp\left(-\frac{1}{2}\big(\xi_\alpha^{(j)}\big)^2\right) = \exp(-\mathcal{S}_k)\,, \tag{S24}$$

with discrete-time action $\mathcal{S}_k \equiv \sum_{j=0}^{k-1} \mathcal{L}^{(j)}$ and Lagrangian

$$\mathcal{L}^{(j)} = \frac{1}{\sigma} \sum_{\alpha} \left(\log \rho_\alpha^{(j+1)}(0) - \log \rho_\alpha^{(j)}(0) - \Delta \log \rho_\alpha^{(j)}\right)^2. \tag{S25}$$

In the small-noise regime, the most likely escape trajectory is the minimum-action path (MAP),

$$\{\log \hat{\boldsymbol{\rho}}^{(0)}, \ldots, \log \hat{\boldsymbol{\rho}}^{(k)}\} \in \arg \min_{\{\log \boldsymbol{\rho}^{(j)}\}} \left\{\mathcal{S}_k \;:\; \rho_\alpha^{(k)} \leq \rho_{\text{extinct}}\right\}. \tag{S26}$$

Among all trajectories that go extinct by cycle $k$, the most likely one (when $\sigma$ is small) is the one that minimizes $\mathcal{S}_k$ subject to the endpoint constraint $\rho_\alpha^{(k)} \leq \rho_{\text{extinct}}$. An example minimum-action path for a particular vector field is shown in Fig. S3(a,b).

#### *S3.B.2. First-passage-time representation of the community lifetime*

With a path defined, the community lifetime is the first cycle at which species $\alpha$ crosses the extinction threshold. Its mean can be written in the standard discrete-time first-passage form as

$$\bar{\tau}_{\text{lifetime}} = \sum_{k \geq 1} k\, \mathbb{P}(\rho_\alpha^{(k)} \leq \rho_{\text{extinct}}). \tag{S27}$$

Using the Onsager–Machlup path weight, the unnormalized weight of trajectories reaching extinction by cycle $k$ is $\propto \exp(-\mathcal{S}_k)$, so the extinction probability is a path integral

$$\mathbb{P}(\rho_\alpha^{(k)} \leq \rho_{\text{extinct}}) \propto \int_{\rho_\alpha^{(k)} \leq \rho_{\text{extinct}}} \exp\big(-\mathcal{S}_k[\log \boldsymbol{\rho}^{(0)}, \ldots, \log \boldsymbol{\rho}^{(k)}]\big) \prod_{j=0}^{k-1} d \log \boldsymbol{\rho}^{(j)}. \tag{S28}$$

In the small-noise regime this integral is dominated by the minimum-action path, giving

$$\mathbb{P}(\rho_\alpha^{(k)} \leq \rho_{\text{extinct}}) \asymp \exp\left(-\frac{V_b(k)}{\sigma}\right), \qquad V_b(k) \equiv \min_{\rho_\alpha^{(k)} \leq \rho_{\text{extinct}}} \sigma \mathcal{S}_k. \tag{S29}$$

Because $\mathcal{S}_k$ contains a factor $1/\sigma$, the combination $\sigma\mathcal{S}_k$ is $\sigma$-independent and plays the role of the Freidlin–Wentzell quasipotential barrier [11]. The sum over $k$ is then dominated by the optimal escape cycle $\hat{k}$ minimizing $V_b(k)$, giving an Arrhenius-type scaling

$$\bar{\tau}_{\text{lifetime}} \asymp A(\sigma) \exp\left(\frac{V_b}{\sigma}\right), \qquad V_b \equiv \min_{k \geq 1} V_b(k), \tag{S30}$$

where $A(\sigma)$ is a subdominant prefactor.

##### S3.B.3. *Building an intuition of action as a quasipotential*

To interpret $V_b$, it is useful to expand the square in the Lagrangian. Define the total observed change in log-density at step $j$ as $d \log \rho_\alpha^{(j)} \equiv \log \rho_\alpha^{(j+1)}(0) - \log \rho_\alpha^{(j)}(0)$, so that the residual is $d \log \rho_\alpha^{(j)} - \Delta \log \rho_\alpha^{(j)}$. Starting from

$$\mathcal{S}_k = \frac{1}{\sigma} \sum_{j=0}^{k-1} \sum_\alpha \left(d \log \rho_\alpha^{(j)} - \Delta \log \rho_\alpha^{(j)}\right)^2,$$

expanding gives

$$\begin{aligned}
\mathcal{S}_k &= \frac{1}{\sigma} \sum_{j=0}^{k-1} \sum_\alpha \left[\left(d \log \rho_\alpha^{(j)}\right)^2 - 2\, d \log \rho_\alpha^{(j)} \cdot \Delta \log \rho_\alpha^{(j)} + \left(\Delta \log \rho_\alpha^{(j)}\right)^2\right] \\
&= -\frac{2}{\sigma} \sum_{j=0}^{k-1} \sum_\alpha d \log \rho_\alpha^{(j)} \cdot \Delta \log \rho_\alpha^{(j)} + \frac{1}{\sigma} \sum_{j=0}^{k-1} \sum_\alpha \left[\left(d \log \rho_\alpha^{(j)}\right)^2 + \left(\Delta \log \rho_\alpha^{(j)}\right)^2\right].
\end{aligned}$$

The first (cross) term is the only contribution that can reduce the action; it accumulates the projection of the observed trajectory onto the deterministic drift. Along the MAP, we approximate this discrete sum by a line integral parameterized by $\log \boldsymbol{\rho}(0)$ along the trajectory:

$$-\frac{2}{\sigma} \sum_{j=0}^{k-1} \sum_\alpha d \log \rho_\alpha^{(j)} \cdot \Delta \log \rho_\alpha^{(j)} \approx -\frac{2}{\sigma} \int_{\text{MAP}} \Delta \log \boldsymbol{\rho} \cdot d \log \boldsymbol{\rho}, \tag{S31}$$

where the line integral is taken along the MAP trajectory in log-density space. For the scalar projected flow shown in Fig. S3(c), this reduces to the integral of $\Delta \log \boldsymbol{\rho}_{\|} \equiv \Delta \log \boldsymbol{\rho} \cdot \boldsymbol{y}$ (the component of the deterministic drift along the unit vector $\boldsymbol{y}$ tanget to the MAP) with respect to arc length along the path. This motivates interpreting the effective barrier $V_b = \sigma \mathcal{S}_{\hat{k}}$ as accumulated motion against the deterministic drift along the minimum-action escape path. The remaining positive contribution,

$$\frac{1}{\sigma} \sum_{j=0}^{k-1} \sum_{\alpha} \left[ \left( d \log \rho_{\alpha}^{(j)} \right)^2 + \left( \Delta \log \rho_{\alpha}^{(j)} \right)^2 \right],$$

acts as a trajectory-dependent prefactor. This recovers the Arrhenius analogy: the lifetime is controlled primarily by an effective barrier $V_b$ associated with the minimum-action escape path (Fig. S3(c)). For $N > 2$, $V_b$ should be interpreted as the accumulated quasipotential along the optimal multidimensional escape trajectory, rather than the height of a one-dimensional potential.

##### *S3.B.4. Linking the escape energy back to invasibility.*

Finally, we connect the escape barrier to the invasibility metric. When $\rho_\alpha \to 0$, the deterministic drift of the focal species approaches the asymptote

$$\Delta \log \rho_{\alpha}^{(j)} \longrightarrow I_{\alpha}(\boldsymbol{\rho}_0^{\setminus \alpha}) + \log \delta, \tag{S32}$$

where $\boldsymbol{\rho}_0^{\setminus \alpha}$ denotes the background composition of the other species (i.e., the community state relevant for evaluating $I_\alpha$). Once the minimum-action escape trajectory enters the low-$\rho_\alpha$ regime (after some cycle $k'$), we may replace $\Delta \log \rho_{\alpha}^{(j)}$ by the constant drift $I_{\alpha}(\boldsymbol{\rho}_0^{\setminus \alpha}) + \log \delta$ for $j \geq k'$. At the level of the action this yields

$$\mathcal{S}_k = \frac{1}{\sigma} \sum_{j=0}^{k-1} \sum_{\alpha} \left( d \log \rho_{\alpha}^{(j)} - \Delta \log \rho_{\alpha}^{(j)} \right)^2, \tag{S33a}$$

$$\approx \frac{1}{\sigma} \sum_{j=0}^{k'-1} \sum_{\alpha} \left( d \log \rho_{\alpha}^{(j)} - \Delta \log \rho_{\alpha}^{(j)} \right)^2 + \frac{1}{\sigma} \sum_{j=k'}^{k-1} \Big( d \log \rho_{\alpha}^{(j)} - I_{\alpha}(\boldsymbol{\rho}_0^{\setminus \alpha}) - \log \delta \Big)^2, \tag{S33b}$$

$$\approx \frac{1}{\sigma} \sum_{j=0}^{k-1} \Big( d \log \rho_{\alpha}^{(j)} - I_{\alpha}(\boldsymbol{\rho}_0^{\setminus \alpha}) - \log \delta \Big)^2, \tag{S33c}$$

where Eqn. (S33b) separates the early part of the trajectory (where the full nonlinear deterministic drift $\Delta \log \rho_{\alpha}^{(j)}$ matters) from the late part (where the low-$\rho_\alpha$ asymptote applies), and Eqn. (S33c) corresponds to the simplifying limit in which the low-$\rho_\alpha$ regime dominates the action.

Under this approximation, the dependence of the action on the background community enters only through $I_\alpha(\boldsymbol{\rho}_0^{\setminus\alpha})$. Consequently, among extinction trajectories satisfying the endpoint constraint $\rho_\alpha^{(k)} \leq \rho_{\text{extinct}}$, the minimum-action escape selects the background composition that minimizes $I_\alpha(\boldsymbol{\rho}_0^{\setminus\alpha})$ (i.e., yields the strongest net drift toward extinction):

$$\hat{\boldsymbol{\rho}}_0^{\setminus\alpha} \in \arg\min_{\boldsymbol{\rho}_0^{\setminus\alpha}} I_\alpha(\boldsymbol{\rho}_0^{\setminus\alpha}). \tag{S34}$$

The same substitution connects the barrier to invasibility. In the low-$\rho_\alpha$ regime, the MAP reduces to a 1D path along the extinction coordinate of species $\alpha$ alone, with the background community frozen at $\hat{\boldsymbol{\rho}}_0^{\setminus\alpha}$. The barrier is therefore a 1D integral along $\log\rho_\alpha$ only:

$$V_b \approx -2\int_{\log\rho_\alpha^*}^{\log\rho_{\text{extinct}}} \Delta\log\rho_\alpha\Big(\log\rho_\alpha;\, \hat{\boldsymbol{\rho}}_0^{\setminus\alpha}\Big)\, d\log\rho_\alpha, \tag{S35}$$

where $\Delta\log\rho_\alpha(\log\rho_\alpha;\, \hat{\boldsymbol{\rho}}_0^{\setminus\alpha})$ is the deterministic drift of species $\alpha$ at log-density $\log\rho_\alpha$ with the background fixed at $\hat{\boldsymbol{\rho}}_0^{\setminus\alpha}$, and $\log\rho_\alpha^*$ is the log-density of species $\alpha$ at the coexistence fixed point. Since the drift is positive (pushing back toward coexistence) and the integration runs downward from $\log\rho_\alpha^*$ to $\log\rho_{\text{extinct}}$, the integral is negative and $V_b > 0$. In the low-$\rho_\alpha$ limit, $\Delta\log\rho_\alpha \approx I_\alpha(\hat{\boldsymbol{\rho}}_0^{\setminus\alpha}) + \log\delta$ is approximately constant, giving

$$\begin{aligned} V_b &\approx -2\left(\log\rho_{\text{extinct}} - \log\rho_\alpha^*\right)\Big(I_\alpha(\hat{\boldsymbol{\rho}}_0^{\setminus\alpha}) + \log\delta\Big) \\ V_b &\mathrel{\substack{\propto\\\sim}} I_\alpha(\hat{\boldsymbol{\rho}}_0^{\setminus\alpha}). \end{aligned} \tag{S36}$$

Thus, the same quantity $I_\alpha(\hat{\boldsymbol{\rho}}_0^{\setminus\alpha})$ that controls invasibility also controls the effective escape barrier and therefore the lifetime through the Arrhenius scaling.

### S4. A PHENOMENOLOGICAL MODEL OF MANY SPECIES

#### S4.A. The model

The phenomenological model assigns each species the general rate equation $\frac{d}{dt}\rho_\alpha = \lambda_\alpha(\rho_{\text{tot}})\rho_\alpha$ (Eqn. 1 in the main text), where the per-capita growth rate $\lambda_\alpha$ is an arbitrary function of total community biomass $\rho_{\text{tot}}$, following the community-state reparameterization of time introduced in [9]. This growth function is discretized into $M$ equal biomass bins of width $\Delta\rho$:

$$\lambda_\alpha(\rho_{\rm tot}) = G_{\alpha,b} \quad \text{if} \quad b\Delta\rho \le \rho_{\rm tot} < (b+1)\Delta\rho, \tag{S37}$$

where $G$ is an $N \times M$ growth rate matrix whose rows index the $N$ species and whose columns index the $M$ biomass bins. A particular community strategy is thus fully encoded in $G$. For small $\Delta\rho$, the species abundances within one cycle are well approximated by the discrete update rule

$$\rho_{\alpha,b+1} = \rho_{\alpha,b} + \frac{G_{\alpha,b}\,\rho_{\alpha,b}}{\sum_\beta G_{\beta,b}\,\rho_{\beta,b}}\Delta\rho. \tag{S38}$$

The dynamics are invariant to the column sums of $G$; matrices are therefore normalized to have unit column sums. The total community biomass is initialized to $\rho_{\rm tot,0} = 1$, and $\Delta\rho$ is held fixed across all bins. After all $M$ bins, i.e. one full growth cycle, the community is diluted by the dilution fraction $\delta = \rho_{\rm tot,0}/\rho_{\rm tot}^{\max} = 1/(1 + M\Delta\rho)$, ensuring the total biomass returns to $\rho_{\rm tot,0} = 1$ at the start of each cycle.

#### S4.B. Calculating co-growth

Co-growth is measured for arbitrary $N$ species as described in SI Sec. S3.

$$I_\alpha \equiv \int_{\delta\rho_{\rm tot}^{\max}}^{\rho_{\rm tot}^{\max}} \frac{\lambda_\alpha(\rho_{\rm tot})}{\sum_{\beta\ne\alpha}\lambda_\beta(\rho_{\rm tot})\rho_\beta(\rho_{\rm tot})} d\rho_{\rm tot}$$

This is implemented numerically in Python as a part of the phenomenological model.

##### *S4.B.1. Numerical calculation of co-growth (minimum invasibility)*

To quantify co-growth, we associate to each species $\alpha$ an invasibility index $I_\alpha$ that measures how easily a rare population of species $\alpha$ can invade a resident subcommunity over a single growth cycle. For a fixed growth-rate matrix $G = \{G_{\beta,b}\}$, we evaluate $I_\alpha(\boldsymbol{\rho}_0^{\setminus\alpha})$ over many choices of subcommunity composition $\boldsymbol{\rho}_0^{\setminus\alpha}$, where for each choice $\rho_{\alpha,0}^{\setminus\alpha}$ is set to a small value and the remaining abundances $\rho_{\beta,0}^{\setminus\alpha} \ge 0$ ($\beta \ne \alpha$) are varied freely; we do not require all $N-1$ other species to be present. Concretely, for a given composition the initial condition is normalized such that $\rho_{\rm tot,0} = 1$, with

$$\rho_{\alpha,0}^{\setminus\alpha} = \frac{\rho_{\rm cutoff}}{N} + \varepsilon_{\rm mach},$$

where we set an extinction cutoff $\rho_{\text{cutoff}} = 10^{-4}$ and $\varepsilon_{\text{mach}}$ is the machine epsilon for the floating-point type used.

We then calculate the generalized invasibility criterion for species $\alpha$ given the initial resident composition $\boldsymbol{\rho}_0^{\setminus\alpha}$ as a discrete sum (an approximation of the continuous integral)

$$I_\alpha(\boldsymbol{\rho}_0^{\setminus\alpha}) = \sum_{b=0}^{M-1} \Delta\rho \, \frac{G_{\alpha,b}}{\sum_{\beta\neq\alpha} G_{\beta,b}\, \rho_{\beta,b} + \varepsilon_{\text{mach}}},$$

where $\Delta\rho$ is the biomass step size per bin. To avoid division by zero, we add a small numerical offset $\varepsilon_{\text{mach}}$ to the denominator.

For each species $\alpha$, we obtain the minimum invasibility ($I_\alpha = \min_{\boldsymbol{\rho}_0^{\setminus\alpha}} I_\alpha(\boldsymbol{\rho}_0^{\setminus\alpha})$) using the default `scipy.optimize.minimize` routine (Python/SciPy), which applies a gradient-based descent (quasi-Newton) method to search over the unconstrained parameterization using a softmax mapping:

$$\tilde{\rho}_\beta = \exp(\rho_\beta - \max_\gamma \rho_\gamma), \quad \beta \neq \alpha$$

$$\rho_{\beta,0}^{\setminus\alpha} = \frac{\tilde{\rho}_\beta}{\sum_{\gamma\neq\alpha} \tilde{\rho}_\gamma}$$

to ensure that values are in the positive orthant, are bounded away from $\infty$, and that the vector is normalized. The optimization starts from uniform composition.

The co-growth value reported in the main text for a given community is then defined as the minimum of these optimized invasibility values,

$$I \equiv \min_{\alpha,\, \boldsymbol{\rho}_0^{\setminus\alpha}} I_\alpha(\boldsymbol{\rho}_0^{\setminus\alpha})$$

which captures the easiest invasion direction into that community and thus quantifies how strongly the community as a whole resists invasion.

#### S4.C. Community lifetime

Starting from the steady-state composition $\boldsymbol{\rho}^*$, we simulate stochastic long-term dynamics by iterating the noisy cycle map. After each growth-dilution cycle, lognormal demographic noise is applied to species abundances as in the main text (Eqn. 4): each species $\alpha$ independently receives a multiplicative perturbation whose logarithm has variance $\sigma/2$, and abundances are renormalized so that $\sum_\alpha \rho_\alpha = 1$ after each step.

$$\rho_\alpha^{(j+1)}(0) = \delta \cdot \rho_\alpha^{(j)}(T) \cdot \frac{1}{Z^{(j)}}\, e^{\sqrt{\sigma/2}\,\xi_\alpha^{(j)}}$$

The noise strength $\sigma$ controls the magnitude of fluctuations; $\sigma = 0$ recovers deterministic dynamics (default: $\sigma = 2.0$).

The community lifetime $\tau_{\text{lifetime}}$ is defined as the index of the first cycle in which any species falls below the extinction threshold. Specifically, a species $\alpha$ is considered extinct at the end of cycle $n$ if

$$\rho^{(n)}_{\alpha,M+1} \cdot \delta \leq \frac{\rho_{\text{cutoff}}}{N}, \tag{S39}$$

with $\rho_{\text{cutoff}} = 10^{-4}$. If no species goes extinct within $n_{\max}$ cycles, the community is assigned $\tau_{\text{lifetime}} = \infty$ (default: $n_{\max} = 10^6$).

To obtain a representative lifetime, we run $R$ independent stochastic replicates from the same $\boldsymbol{\rho}^*$, each with independently drawn noise sequences (default: $R = 100$). The mean lifetime $\bar{\tau}_{\text{lifetime}}$ over replicates with finite $\tau_{\text{lifetime}}$ is the quantity reported in the main text.

### S4.D. Ensemble construction

#### *S4.D.1. Base ensemble construction*

We initialize an ensemble of communities with stable coexistence using random growth matrices $G_{\alpha,b}$ drawn from an exponential distribution and normalized so that $\sum_\alpha G_{\alpha,b} = 1$ for each biomass bin $b$. The matrix is initialized to have a larger species pool ($N_{\text{pool}} > N$) than the eventual number of species that we want to keep, to increase the probability of any given growth matrix producing a coexisting sub-community of size $N$. Each community is then iterated through serial dilution cycles from a uniform initial composition (default: 10,000 warmup cycles), keeping only the communities in which exactly $N$ species remain above the extinction threshold ($\rho_\alpha > \rho_{\text{cutoff}}/N$). Each resulting matrix is restricted to the rows of its coexisting species, and the columns are rescaled to restore unit column sums.

For each community in the filtered ensemble, the steady-state composition $\boldsymbol{\rho}^*$ is then computed by iterating additional growth-dilution cycles from a uniform initial composition until convergence. Convergence is declared when the change in all species abundances over one further cycle satisfies $|\rho_\alpha - F(\boldsymbol{\rho})_\alpha| \leq \rho_{\text{cutoff}}/N$ for all $\alpha$, where $F$ denotes the deterministic (noiseless) cycle map. Communities that fail to converge or have further extinction events (i.e., any species falls below the extinction threshold) are discarded. The resulting $\boldsymbol{\rho}^*$ is the starting composition used for lifetime simulations (SI Sec. S4.C).

*S4.D.2. Replica-exchange Monte Carlo*

Starting from this coexistence-enforcing initial ensemble $G^{(c)}$ ($c$ denotes the community number), we then optimize each community by REMC using co-growth as the sole objective. The co-growth of community $c$ is

$$I^{(c)} \equiv \min_{\alpha,\, \boldsymbol{\rho}_0^{\setminus\alpha}} I_\alpha^{(c)}(\boldsymbol{\rho}_0^{\setminus\alpha})$$

as described in the previous subsection (SI Sec. S4.B). In practice, to avoid repeatedly evaluating co-growth for matrices that clearly do not coexist, we first run a shorter serial dilution simulation (100 cycles) from a uniform initial composition and count the number of species whose abundance exceeds an extinction threshold ($\rho_\alpha > \rho_{\text{cutoff}}/N$, $\rho_{\text{cutoff}} = 10^{-4}$). If this number is less than the total species number, we set $I^{(c)} = 0$ and do not perform the full co-growth calculation. Only matrices that may support full coexistence are passed to the full co-growth routine.

Elementary Monte Carlo moves act on the growth matrices via a mutation operator. For each community and Monte Carlo step, we select $n_{\text{mut}} = 1$ entries of the matrix $G^{(c)}_{\alpha,b}$ uniformly at random over species–bin pairs $(\alpha, b)$. The selected entries are replaced by new values sampled independently from an exponential distribution with mean 1. After mutation, each column (i.e., each bin $b$ for a fixed community $c$) is renormalized

$$\sum_\alpha G^{(c)}_{\alpha,b} = 1,$$

to ensure that the overall scale of growth rates per bin remains fixed while the allocation across species is reshuffled.

We run REMC with $N_{\text{rep}} = 10$ temperature replicas, arranged on a logarithmic ladder

$$T_r = T_{\min}\left(\frac{T_{\max}}{T_{\min}}\right)^{\frac{r-1}{N_{\text{rep}}-1}}, \quad r = 1, \ldots, N_{\text{rep}},$$

with $T_{\min} = 0.1$ and $T_{\max} = 10$ (inverse temperatures $\beta_r = 1/T_r$). Each replica performs $N_{\text{MC}} = 2500$ Monte Carlo mutation–selection steps at its assigned temperature. Within each replica, a proposed mutation to co-growth value $I'$ is accepted with the Metropolis probability

$$p_{\text{acc}} = \min\bigl(1,\, \exp[(I' - I)/T_r]\bigr),$$

where $I$ is the current co-growth of the community. After every 100 local steps, exchange moves are attempted between each pair of neighboring replicas $(r, r+1)$ with acceptance probability

$$p_{\text{swap}} = \min\bigl(1,\, \exp[(\beta_r - \beta_{r+1})(I_{r+1} - I_r)]\bigr),$$

where $I_r$ and $I_{r+1}$ are the co-growth values in replicas $r$ and $r+1$.

At the end of the REMC run, each replica contains an ensemble of optimized matrices. We then perform a quality-control step on each replica separately: we (a) remove duplicate communities by retaining only unique growth matrices along the community axis, and (b) re-check coexistence by computing steady-state abundances and discarding any matrices that do not support full coexistence of all species. For the remaining matrices, any missing fitness values are recomputed using the invasibility procedure. The final ensemble used in the analysis is obtained by consolidating the coexistence-verified, unique matrices from all replicas.

#### S4.E. Arrested development upon removal of a single species

To quantify how individual species contribute to the pace of community development within a growth cycle, we compare the time required for a community to traverse all biomass bins with and without each species present ("leave-one-out," LOO). For a given community with steady-state composition $\boldsymbol{\rho}^*$ and growth-rate matrix $G_{\alpha,b}$, we compute the time increment associated with biomass bin $b$ as

$$\Delta t_b = \frac{\Delta\rho}{\sum_\alpha \rho_{\alpha,b}\, G_{\alpha,b}}, \tag{S40}$$

where $\rho_{\alpha,b}$ is the abundance of species $\alpha$ at the beginning of bin $b$, and $\Delta\rho$ is the fixed biomass increment per bin. The total time to complete a growth cycle is then

$$T_{\text{full}} = \sum_{b=0}^{M-1} \Delta t_b = \sum_{b=0}^{M-1} \frac{\Delta\rho}{\sum_\alpha \rho_{\alpha,b}\, G_{\alpha,b}}. \tag{S41}$$

To assess "arrested development" upon removal of species $\alpha$, we construct a modified initial condition in which species $\alpha$ is deleted from the steady-state composition,

$$\rho^{\setminus\alpha}_{\alpha,0} = 0, \qquad \rho^{\setminus\alpha}_{\beta,0} = \rho^*_{\beta,0} \quad (\beta \neq \alpha), \tag{S42}$$

and evolve this altered community through one growth cycle using the same within-cycle dynamics. This yields a new sequence of abundances $\rho^{\setminus\alpha}_{\beta,b}$, from which we compute the corresponding bin-wise time increments,

$$\Delta t_b^{\setminus\alpha} = \frac{\Delta\rho}{\sum_\beta \rho^{\setminus\alpha}_{\beta,b}\, G_{\beta,b}}, \tag{S43}$$

and the total LOO cycle time

$$T^{\setminus\alpha}_{\mathrm{LOO}} = \sum_{b=0}^{M-1} \Delta t_b^{\setminus\alpha}. \tag{S44}$$

For each community, we repeat this procedure for every species $\alpha = 1, \ldots, N$, obtaining a set of LOO times $\{T^{\setminus\alpha}_{\mathrm{LOO}}\}$. We then form the dimensionless ratio

$$\frac{T^{\setminus\alpha}_{\mathrm{LOO}}}{T_{\mathrm{full}}} \geq 1, \tag{S45}$$

which measures how much the removal of species $\beta$ slows the progression through a growth cycle. In the main text we report, for each community, the maximal slow-down across all single-species deletions,

$$\max_\alpha \frac{T^{\setminus\alpha}_{\mathrm{LOO}}}{T_{\mathrm{full}}}, \tag{S46}$$

and plot this quantity as a function of co-growth. Communities with high co-growth show larger values of this ratio, indicating a stronger dependence on all member species for maintaining rapid development through the growth cycle.

---

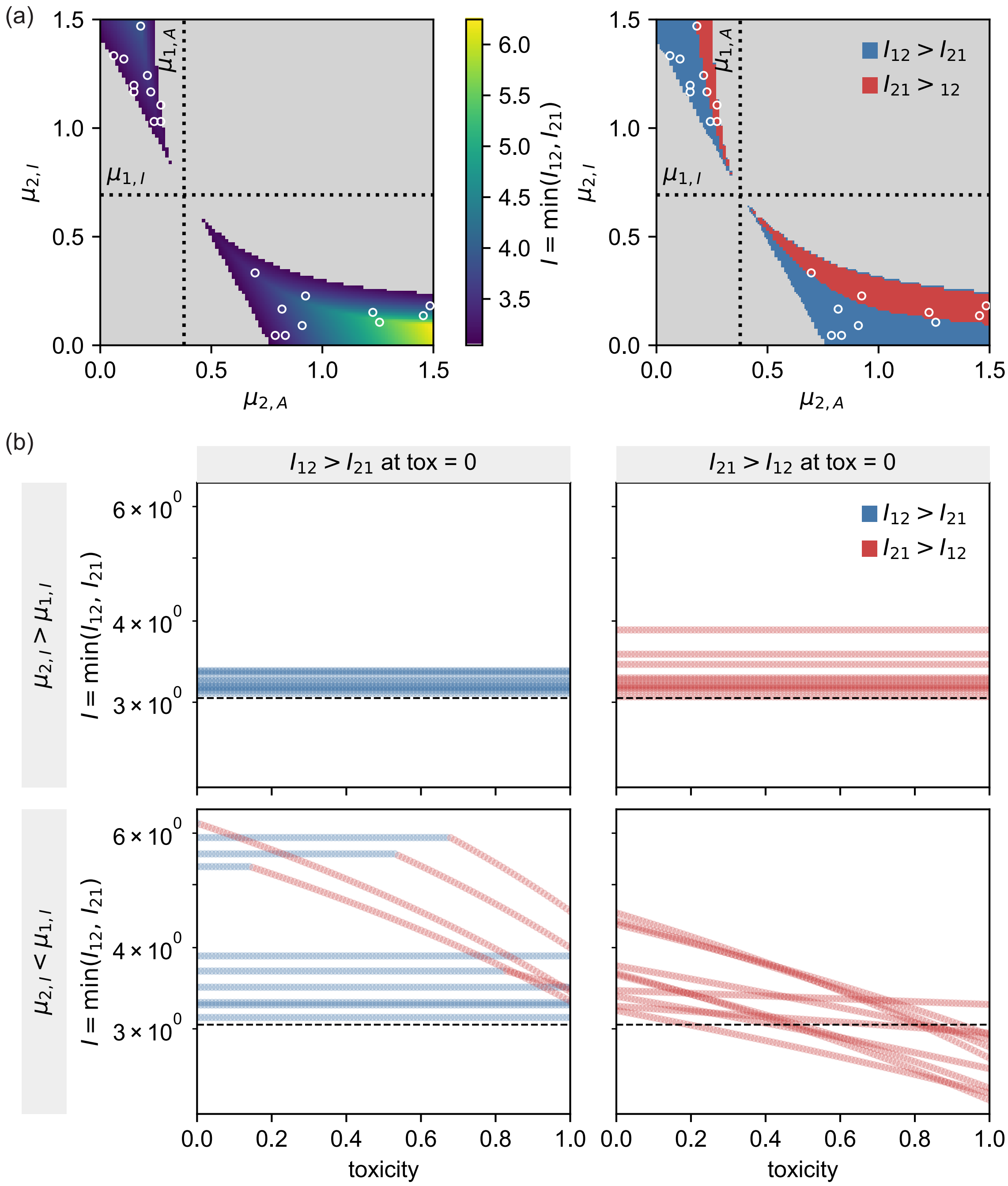


FIG. S2. **Parameter space exploration and toxicity sweep for Anchor 02 ($\mu_{1,A} = 0.377$, $\mu_{1,I} = 0.692$).** Same layout as Fig. S1. This anchor was chosen as a contrast to Anchor 01: it has a similar overall coexistence structure but different effect of toxicity on co-growth.

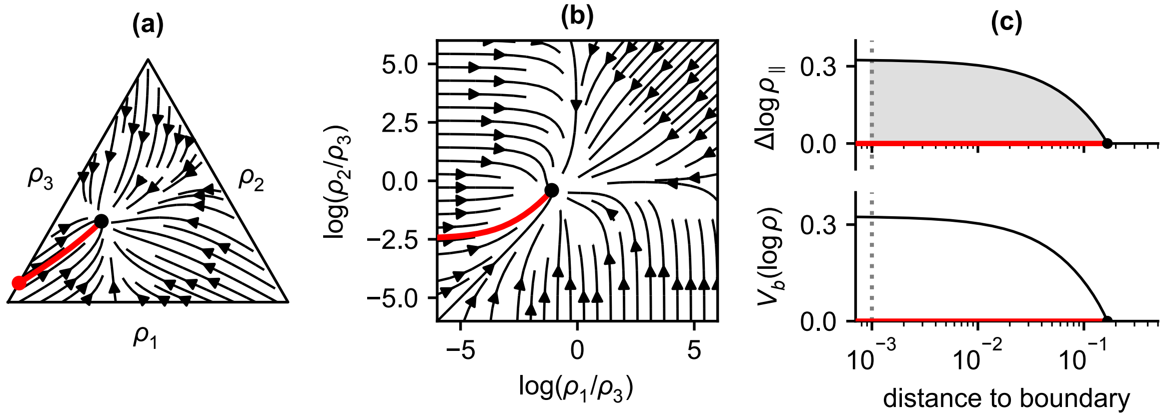


FIG. S3. **Community lifetime in a multi-species model is linked to the quasipotential along the minimum action path.** **(a)** Deterministic change in community composition over growth cycles, $\Delta \log \boldsymbol{\rho}$, shown as a stream plot on the simplex defined by $\rho_1 + \rho_2 + \rho_3 = \rho_{\text{tot}}$. In this plot, we set $\rho_{\text{tot}} = 1$. The black circle denotes the stable fixed point. The red curve indicates the minimum action path (MAP), which minimizes the Freidlin–Wentzell action and represents the most probable escape trajectory from the stable state to the extinction boundary ($\rho_1 = 0$, $\rho_2 = 0$, or $\rho_3 = 0$). **(b)** The same flow field represented in logarithmic coordinates, $\log(\rho_1/\rho_3)$ and $\log(\rho_2/\rho_3)$. **(c)** *Top:* Projection of the vector field along the MAP as a function of distance from the extinction boundary, $\Delta \log \boldsymbol{\rho}_{\|}$. Dotted lines indicate the extinction threshold. *Bottom:* The quasipotential $V_b$, defined as the line integral of the projected flow along the MAP.
